# Boreal moss-microbe interactions are revealed through metagenome assembly of novel bacterial species

**DOI:** 10.1101/2023.04.06.535926

**Authors:** Sarah Ishak, Jonathan Rondeau-Leclaire, Maria Faticov, Sébastien Roy, Isabelle Laforest-Lapointe

## Abstract

Moss-microbe interactions contribute to ecosystem processes in boreal forests. Yet, how host-specific characteristics and the environment drive the composition and metabolic potential of moss microbiomes is still poorly understood. In this study, we use shotgun metagenomics to identify the taxonomy and metabolic potential of the bacteria of four moss species of the boreal forests of Northern Québec, Canada. To characterize moss bacterial community composition and diversity, we assembled the genomes of 110 potentially novel bacterial species. Our results highlight that moss genus, species, gametophyte section, and to a lesser extent soil pH and soil temperature, drive moss-associated bacterial community composition and diversity. In the brown gametophyte section, two *Stigonema* spp. showed partial pathway completeness for photosynthesis and nitrogen fixation, while all brown-associated Hyphomicrobiales had complete assimilatory nitrate reduction pathways and many nearly complete carbon fixation pathways. Several brown-associated species showed partial to complete pathways for coenzyme M and F420 biosynthesis, important for methane metabolism. In addition, green-associated Hyphomicrobiales (*Methylobacteria* spp.) displayed potential for the anoxygenic photosystem II pathway. Overall, our findings demonstrate how host-specific characteristics and environmental factors shape the composition and metabolic potential of moss bacteria, highlighting their roles in carbon fixation, nitrogen cycling, and methane metabolism in boreal forests.

## Introduction

Boreal forests are one of Earth’s largest biomes, covering 29% of land surfaces [1] and storing at least 30% of global carbon [2]. Within this ecosystem, a diverse understory vegetation thrives, consisting of vascular plants, lichens, and bryophytes (small non-vascular plants including liverworts, hornworts, and mosses). Ubiquitous in boreal forests, mosses are crucial contributors to key ecosystem processes, notably to nitrogen (N) fixation and carbon (C) sequestration [3], which are also facilitated by moss symbiotic microorganisms [4, 5]. Moss microbes contribute around 50% of N inputs to boreal forests, play crucial roles in facilitating C sequestration in peatlands, and facilitate host adaptation in degraded landscapes [6]. Moss-associated microbial communities are notably dominated by bacteria from the Proteobacteria, Acidobacteria, Actinobacteria, and Cyanobacteria phyla [7–9]. Driven by the pressures of accelerating climate change, boreal ecosystems face a temperature increase of ∼2.5°C before the end of the century, which could alter moss-microbe interactions with major implications for global N [10] and C cycles [11]. In addition, boreal forests are often harvested intensively for forestry or disturbed by mining activities [12, 13]. Considering the strong pressures of climate change on boreal forests combined with the ever-increasing anthropogenic impacts on land use, improving our understanding of the drivers and functions of moss-microbe interactions is imperative to mitigate the risks of altering this precious habitat [14, 15].

Moss microbial communities can be affected by many local abiotic and biotic factors [8, 16]. For example, soil pH and soil temperature have been shown to structure bacterial communities associated with *Sphagnum* moss species [9, 17, 18]. Phylogenetic relationships and host species are also important drivers of moss bacterial community diversity and community composition [5, 9, 20]. While different moss species have been shown to associate with contrasting bacterial communities, closely related moss species are likely to harbor more similar microbial taxa than more distantly related ones [18, 21, 22]. This convergence can be attributed to shared host physiological traits and ecological niches. For example, Holland-Moritz *et al*. [23], showed that phylogenetic relationship between *Sphagnum* species explained up to 65% of the variation in bacterial community composition. Interestingly, moss species identity also influences the effects of environmental factors on bacterial diversity and community composition [18]. While previous studies have often focused on characterizing the microbial taxonomy and functions of dominant boreal mosses such as *Pleurozium schreberi* [24] and peat mosses of the genus *Sphagnum* [25], scant research is available on other moss genera such as *Polytrichum* and *Dicranum* (but see [26]), which are common in Canadian boreal forest [27]. Importantly, these taxa are key candidates for remediation and restoration efforts as they often thrive in degraded landscapes [28]. Thus, there is an important knowledge gap on moss-microbe interactions beyond common moss genera. Considering the urgency to develop conservation strategies for bryophytes, it is crucial to deepen our understanding of moss-microbe interactions and their functional roles in unaltered boreal ecosystems by investigating understudied moss species, their phylogenetic relationships, and local environmental factors.

Beyond moss species identity, physiological differences within a single host, such as disparities in tissue types along the moss gametophyte, could also influence moss-microbe interactions. Gametophytes are the dominant life stage of mosses, separated along their vertical gradient in two distinct sections: a living “green” section and an underlying, decaying “brown” section [29]. These sections can differ in microbial diversity, as well as in the composition and functional potential of their microbial communities. [30, 31]. The green section can harbor photosynthetically active microbes [32, 33] and is associated with N-fixing cyanobacteria [34]; it is often the focus of studies on moss-bacteria associations. However, the brown section is much less studied even though it is thought to be an important source of nutrients for specific groups of microbes, such as bacteria [33, 35]. Notably, green and brown sections of the moss gametophyte are exposed to distinct microenvironmental conditions [36], whereby variations in light, soil pH, moisture, and temperature can impact moss microbial diversity and functions. Overall, to gain a detailed overview of moss-microbe interactions, it is necessary to measure how gametophyte sections assemble divergent microbial communities and functional profiles, and whether these differences are consistent among moss species as well as along environmental gradients.

In recent years, metagenome-resolved community profiling became a widespread approach to estimate environmental microbial composition and functions [37]. The advent of whole metagenome sequencing (WMS) revolutionized the field of microbial ecology, making it possible to assemble metagenome-assembled genomes (MAGs) and thus delve into the so-called “dark matter” of microbiomes [38]. By leveraging the entire genomic material of a sample, WMS not only enables the discovery of novel or uncultured microbial species; it also provides a more complete picture of the metabolic potential of community members [39]. WMS is therefore particularly useful in characterizing understudied microbiomes, such as that of boreal mosses. The discovery of novel bacterial species, along with an improved understanding of their metabolic potential in different gametophyte sections and environmental conditions, is important for predicting how these interactions will shift as climate change and human activities alter boreal ecosystems.

Here, we measure the influence of moss identity, gametophyte section, and abiotic environment on moss bacterial communities and their metabolic potential. We focus on four boreal moss species in the Eeyou Istchee (James Bay) region of Québec, Canada: *Polytrichum juniperinum*, *P. commune*, *P. piliferum*, and *Dicranum undulatum* (Fig. 1ab). We use WMS and a combination of cutting-edge bioinformatic tools to (i) characterize moss-microbe interactions, (ii) assemble metagenomes, and (iii) identify novel bacterial species associated with boreal mosses. Our objectives are to (1) characterize the diversity, community composition, and metabolic potential of moss-associated bacteria; and (2) quantify the relative importance of moss species, gametophyte section, soil pH, and soil temperature as drivers of moss-bacteria interactions.

**Figure 1.**
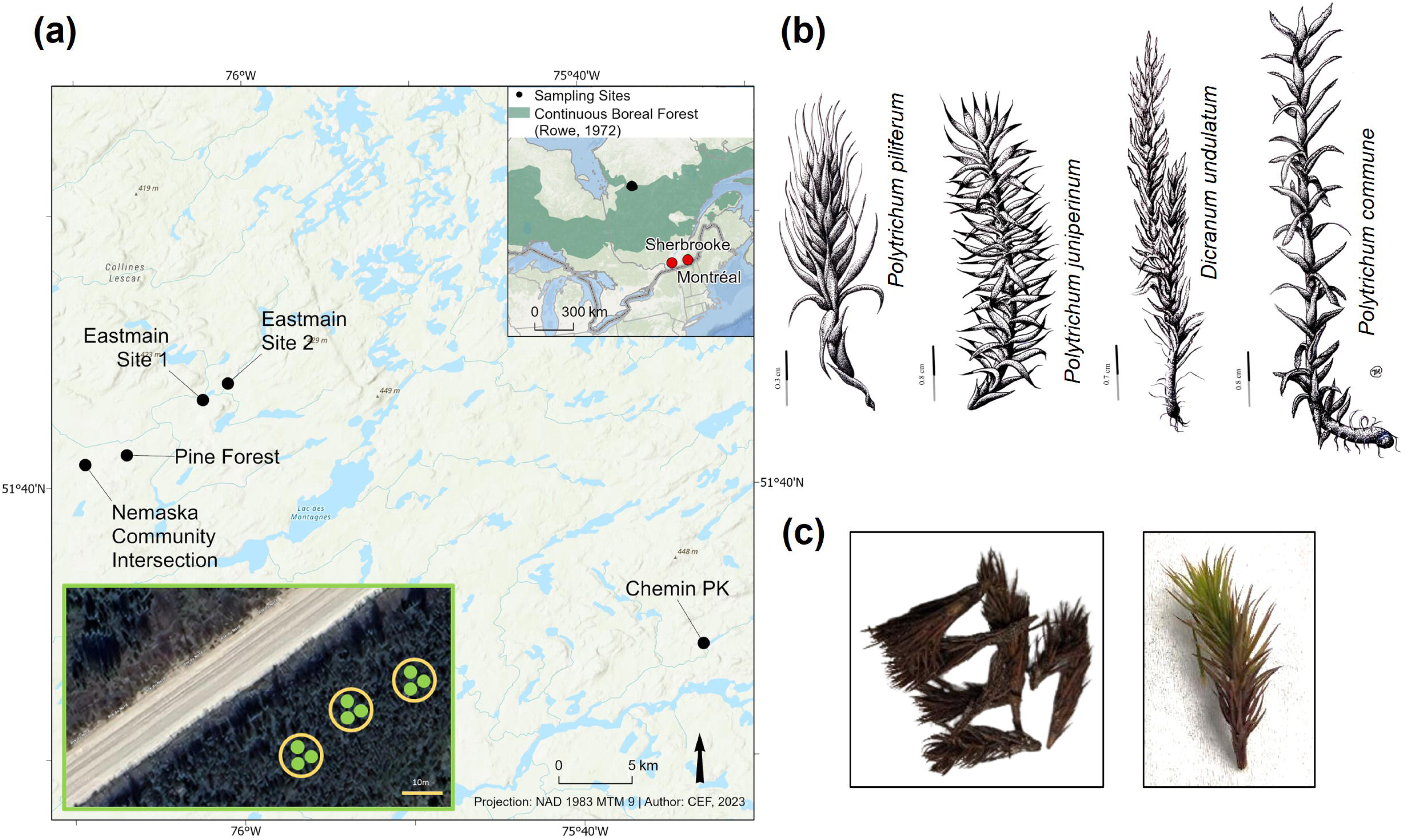
Sampling design, location, and collected moss species. Panel (a) shows the sampling layout: at each location, three microsites were chosen at random (indicated by yellow circles). Within each microsite, three replicate samples were collected (shown as green circles). Inset in the top right of Panel (a) shows a map of Québec with the sampling sites located within the continuous boreal forest region. Panel (b) shows the illustrations of each of four moss species (drawings were created by Isabel Ramirez). Panel (c) shows the images of green and brown sections of the moss gametophyte.

## Results and Discussion

### Bacterial community composition and diversity are driven by moss identity, gametophyte section, soil pH, and soil temperature

First, we investigated the influence of moss genus, moss species (nested within moss genus), gametophyte section, soil pH, and soil temperature on bacterial community composition and diversity (Table 1a). We then narrowed our focus on the three *Polytrichum* spp. (excluding *Dicranum undulatum*; Table 1b). This sequential approach allowed us to disentangle whether closely related moss species are likely to harbor more similar microbial taxa than more distantly related ones. Both moss genus and species (nested within genera) were strong drivers of bacterial community composition (Genera: R^2^ = 13.8%, p-value = 0.001; Species: R^2^ = 12.3%, p-value = 0.001; Fig. 2a, Table 1a). Notably, when restricting the analysis to the three *Polytrichum* spp., the effect of moss species on bacterial community composition remained significant (R^2^ = 19.7%, p-value = 0.001; Table 1b). Thus, our results highlight that differences in bacterial community composition are driven by both moss genus and, within these genera, by moss species. These results are in line with previous studies in which moss phylogeny was shown to be an important driver of bacterial community composition [4, 8, 25]. This finding illustrates a potential co-evolutionary relationship between mosses and some of the bacteria members within the community [17, 40]. Yet, more studies are needed to fully understand the nature and mechanisms driving these genus- and species-specific interactions. We also found that bacterial community composition differed between green and brown gametophyte sections, regardless of whether *D. undulatum* samples were included in the model or not (R^2^ = 12.7%, p-value = 0.001; R^2^ = 17.2%, p-value = 0.001, respectively; Fig. 2b, Table 1ab). This variation between sections can be explained by differences in physiology [41] and micro-environmental conditions (e.g., soil pH, moisture, temperature) [36]. In addition, we observed a significant, albeit small, interaction between gametophyte section and moss genus (R^2^ = 2.6%, p-value = 0.001; Table 1a), as well as moss species (R^2^ = 1.3%, p-value = 0.042; Table 1b), highlighting that the effect of gametophyte section on bacterial community composition differs among moss species. In both models, whether *D. undulatum* was included or not, we found that soil temperature (with *Dicranum:* R^2^ = 7.4%, p-value = 0.001; without *Dicranum*: R^2^ = 6.9, p-value = 0.001) and pH (R^2^ = 5.1%, p-value = 0.001 and R^2^ = 4.1%, p-value = 0.001 respectively) were significant drivers of bacterial community composition. Interestingly, the effect of abiotic factors on community composition was weaker than that of moss genus and species (Table 1). Finally, the interaction of soil pH and temperature affected moss bacterial community composition (Table 1), in coherence with the important roles of abiotic factors in structuring moss-microbe interactions across moss species [9, 17, 18].

**Figure 2.**
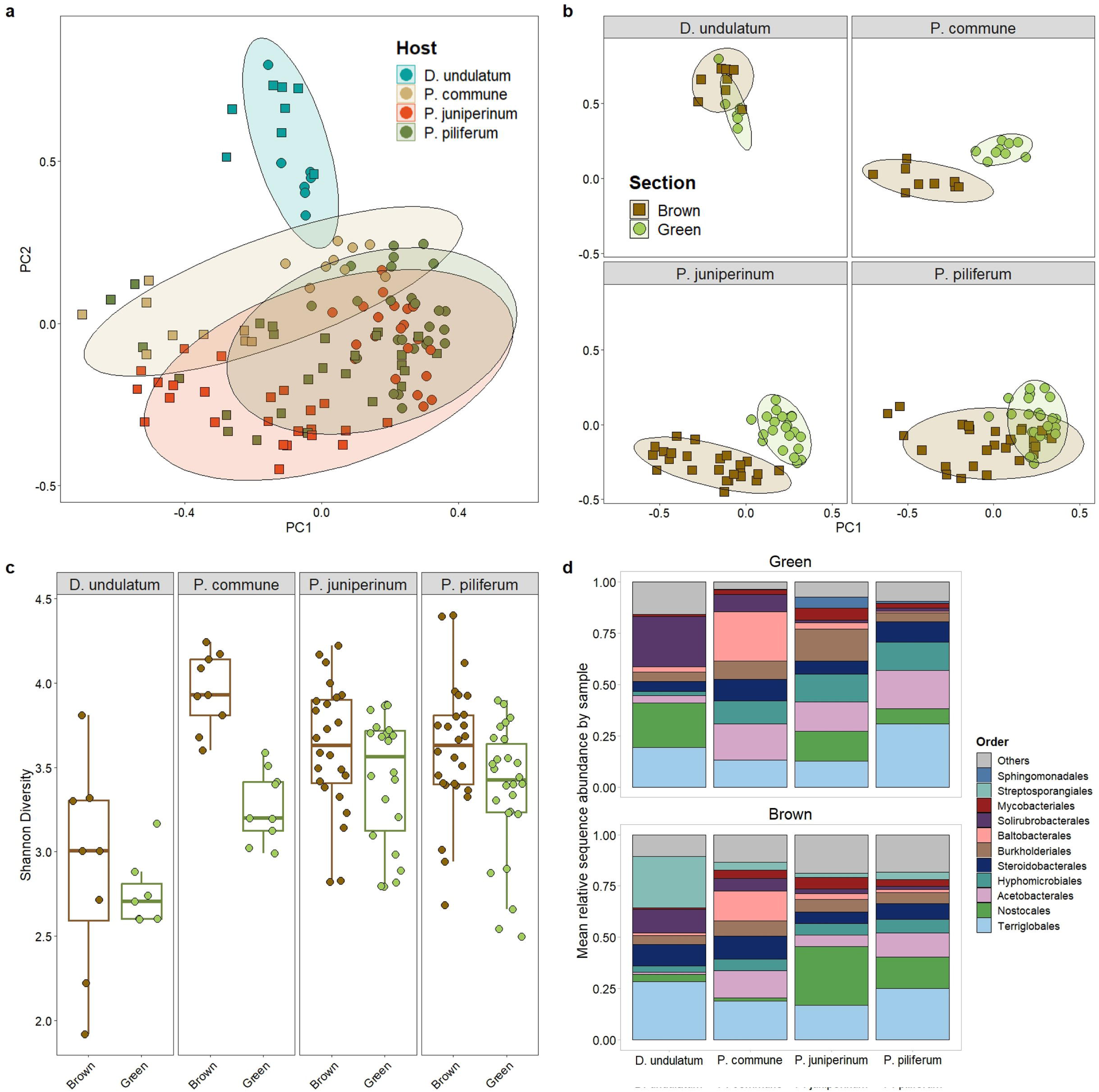
(a, b) Two-dimensional principal coordinates analysis (PCoA) of bacterial communities based on Bray-Curtis dissimilarity index, calculated using variance-stabilizing-transformed abundance table. Each point represents one moss sample, and ellipses indicate a 95% bivariate confidence interval. Each ellipse colour represents a moss species (a) or a moss gametophyte section (b). (c) Differences in alpha diversity (Shannon diversity) of moss bacterial communities. Showed are four moss species: *P. juniperinum* (N_brown_ = 24, N_green_ = 22), *P. commune* (N_brown_ = 9, N_green_ = 9), *P. piliferum* (N_brown_ = 26, N_green_ = 26), and *D. undulatum* (N_brown_ = 8, N_green_ = 7). (d) Mean sequence relative abundance by bacterial orders between green (top panel) and brown (bottom panel) gametophyte sections across four moss species.

**Table 1.**
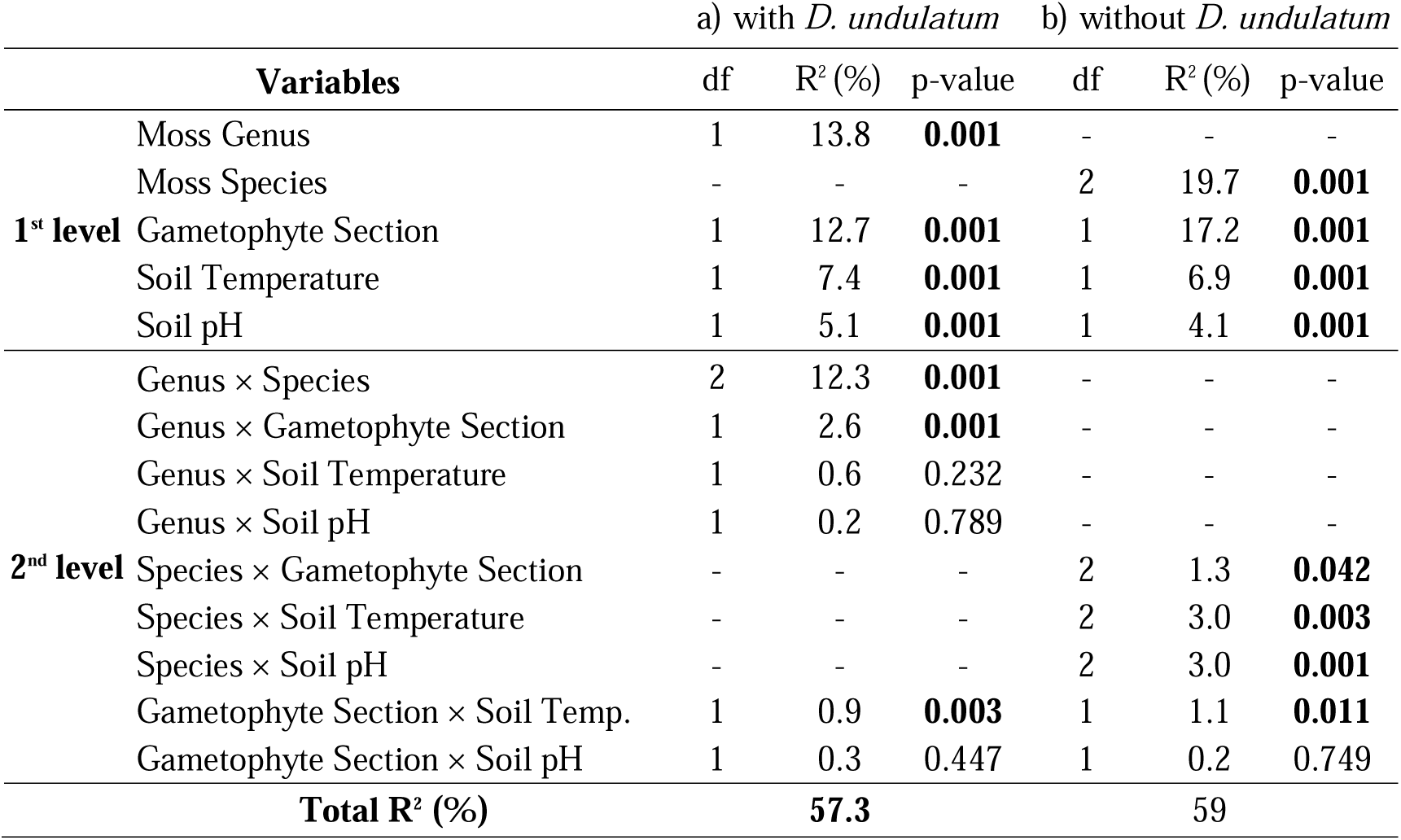
Moss genus, moss species nested within moss genus, gametophyte section, soil pH, soil temperature, and their interactions drive moss bacterial community composition. Shown are the PERMANOVA on Bray-Curtis dissimilarity (n = 131 and n = 121 samples respectively) degrees of freedom (df), determination coefficients (R^2^), and p-values (significant values are shown in bold).

Bacterial alpha diversity differed between moss genera, with alpha diversity (bacterial Shannon diversity) being higher for *Polytrichum* than for *D. undulatum* (χ*^2^* = 8.6, p-value = 0.003; Table 2a, Fig. 2c). However, we did not detect a significant association between moss species and bacterial alpha diversity, neither when nesting species within moss genera (χ*^2^ =* 0.07, p-value = 0.97; Table 2a, Fig. 2c) nor when narrowing the model to *Polytrichum* spp. (χ*^2^ =* 0.08, p-value = 0.96; Table 2b, Fig. 2c). Of note, alpha diversity in the green section was consistently lower than in the brown section of the gametophyte. This pattern was consistent between moss genera (χ*^2^* = 29.2, p-value < 0.001; Fig. 2c, Table 2a) and among *Polytrichum* spp. (χ*^2^* = 35.7, p-value < 0.001; Fig. 2c, Table 2b). This could be explained by (i) a phenomenon of niche dominance of a few taxa in the green section or by (ii) the proximity of the brown section to the soil, which is known to be extremely rich in microorganisms. As such, the brown section could behave as an interface in which both soil and moss-associated bacteria can thrive. This section is also mostly shaded from UV and may contain less chlorophyll than the above active, green section [42]. Finally, the underlying brown section tends to have higher moisture contents due to the boundary layer effect (i.e., the physical phenomenon in which wind velocity is reduced); an effect that minimizes moisture loss [43]. Among abiotic factors, soil temperature was the only factor showing a significant association with moss bacterial alpha diversity, evident in both models, with and without *D. undulatum* (χ*^2^* = 6.3, p-value = 0.01 and χ*^2^* = 6.6, p-value = 0.01; Fig. 2c, Table 2ab). Furthermore, we did not detect a significant effect of the interaction between gametophyte section and moss genus or species on bacterial alpha diversity (Table 2).

**Table 2.**
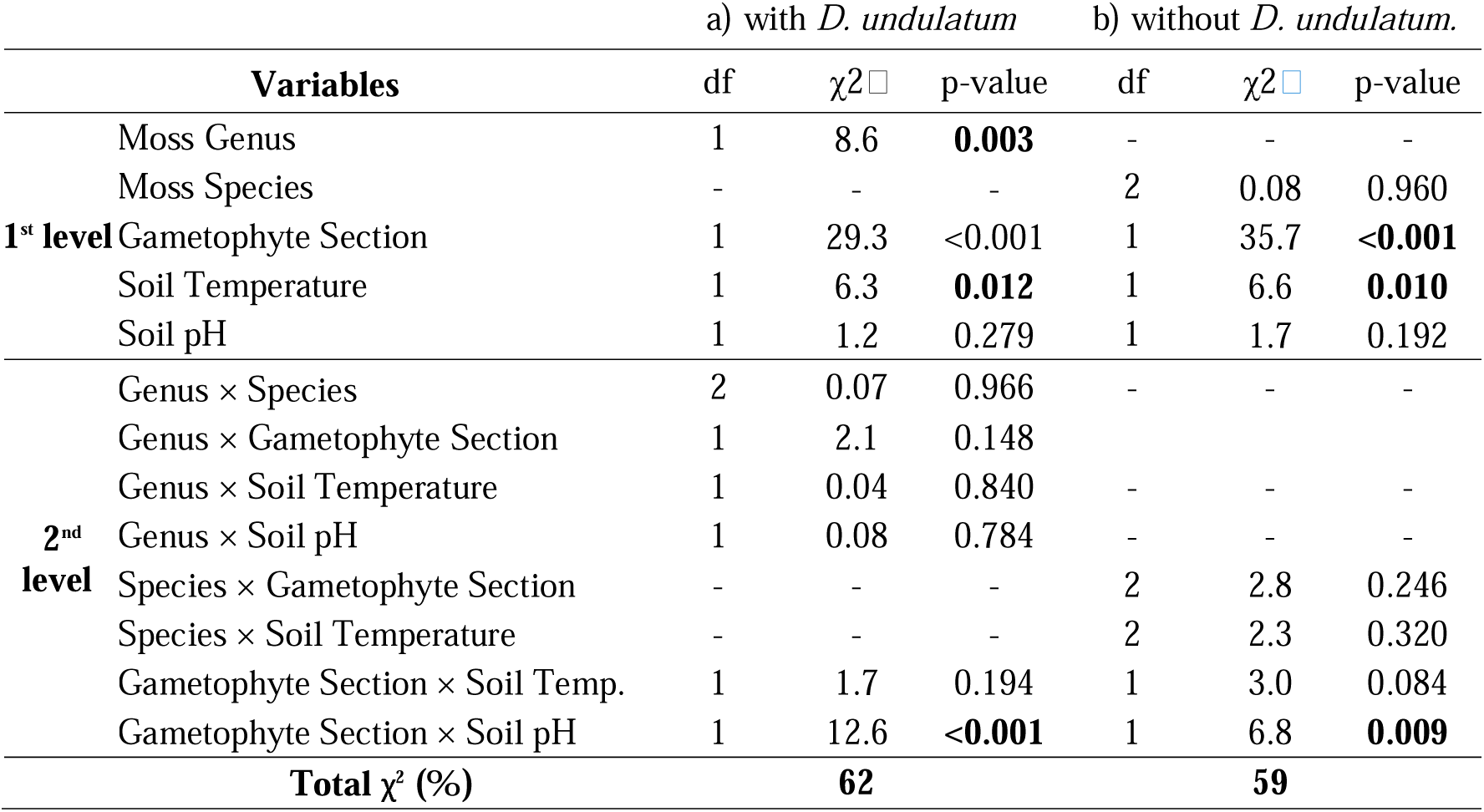
The effects of moss genus, moss species nested within moss genus, gametophyte section, soil pH, and temperature on moss bacterial alpha-diversity (Shannon index). Shown are the linear mixed-effects model (n = 131 and n = 121 samples respectively) test statistics (χ*^2^*), degrees of freedom (df), and p-values (significant values are shown in bold).

In summary, these results show that moss genus, gametophyte section, soil pH, and soil temperature are important drivers of moss bacterial community composition and diversity, but that moss species only affects the former. Our work further demonstrates that green and brown sections within a moss gametophyte differ in bacterial community composition (Fig. 2d) and alpha diversity, emphasizing the relationship between moss characteristics and the associated bacterial communities.

### Metagenome assembly reveals that novel bacteria constitute most of moss bacterial community

By assembling metagenomes, we discovered 110 potentially novel bacterial species whose genomes were not represented in reference databases. Specifically, we performed metagenome co-assembly, binning, and dereplication, which produced 157 non-redundant species-level MAGs. We filtered out MAGs with quality scores (QS) ≤ 0.50 (QS = completeness - 5 × contamination), yielding a subset of 118 MAGs of medium to high quality (82.46 ± 11.68 % completeness, 1.97 ± 1.26 % contamination). In order to estimate which MAGs belong to undescribed species, we compared them to every known representative bacterial genome using a combination of tools based on mash distance and average nucleotide identity (see *Methods*). This resulted in a set of draft genomes from 110 putative novel bacterial species spanning 23 orders across nine phyla. Hereafter, we refer to this set as the novel species-level MAGs (nMAGs), in that they constitute genomes that are non-redundant with any published reference genome at the species level. These nMAGs have a mean length of 4.0 ± 1.2 Mbp, QS of 0.73 ± 0.13, and L50 of 11.9 kb ± 10.5. Full quality statistics of the nMAGs, including the number of contigs and proportion of GC content, are reported in Supplementary Data 1 and summarized in Fig. 3.

**Figure 3.**
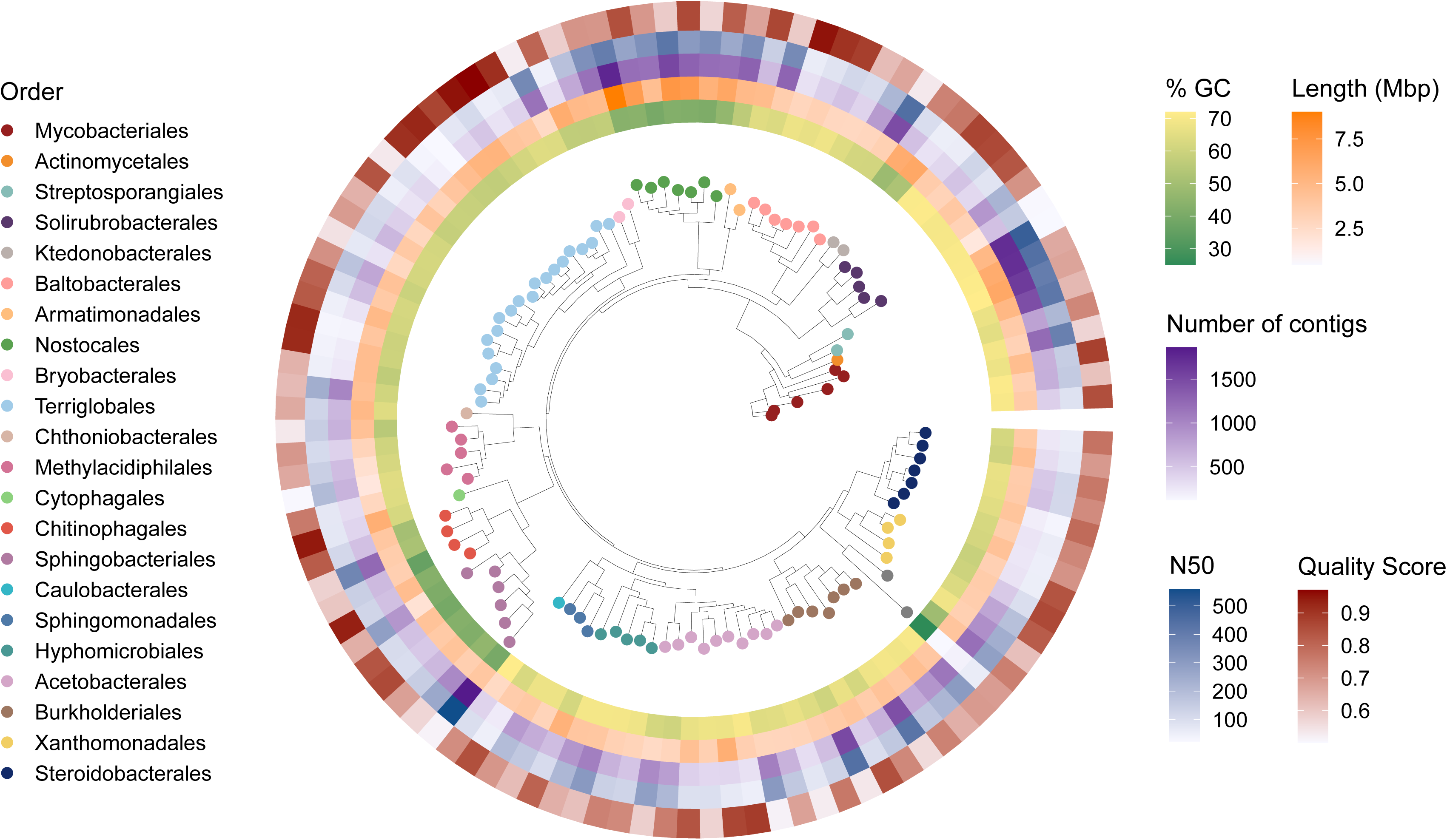
Novel species-level metagenome-assembled genomes obtained with co-assembly. From inner to outer layers: proportion of guanine and cytidine in assembly nucleotides (%LJGC), genome size (length, Mbp), number of contigs in each MAG, length of the shortest of the largest contigs required to cover 50% of the total assembly length (N50) and quality score (QS). The inner section shows the phylogenetic tree of 110 metagenome-assembled genomes constructed using PhyloPhlAn and built phylogenetic trees using the ggtree package, allowing a cross-validation with their assigned taxonomy in the GTDB database. Two clades were renamed using NCBI taxonomy, as the GTDB nomenclature used illegitimate names: Nostocales (Cyanobacteriales) and Hyphomicrobiales (Rhizobiales). The different colored dots on the branches of the tree correspond to the colors of the bacterial orders listed in the legend.

To gain a more detailed understanding of the microbial communities of boreal mosses, we added the 110 nMAGs to the 85,205-reference genome GTDB used for community profiling, resulting in an average 4.3-fold increase in sample containment (a measure of proportion of sample *k*-mers included in the minimal reference genomes set that encompasses all k-mers in a sample; see Methods; Supplementary Fig. S1). This increase demonstrates the effectiveness of using *de novo* metagenome assembly to identify and describe previously unknown microbial species, which is crucial for environmental datasets, and to increase the proportion of sample information to perform ecological analyses. As detailed in Methods, all results presented here (including the above section on diversity) rely on community composition estimation performed using the improved reference database.

Of note, we modified the GTDB taxonomy used to annotate our genomes according to the latest nomenclature in the *List of Prokaryotic names with Standing in Nomenclature* (LPSN) [44]. Specifically, GTDB classifies Hyphomicrobiales (LPSN nomenclature) as Rhizobiales, so we corrected our annotations to use Hyphomicrobiales throughout. GTDB also uses the order Cyanobacteriales, which encompasses species of many different orders (e.g., Oscillatoriales, Synechococcales, Chroococcales, etc.) according to LPSN. Since every member of the class Cyanobacteria found in our samples belonged to the Nostocaceae family, we renamed the Cyanobacteriales order to Nostocales, following LPSN.

We recovered seven MAGs belonging to the order Baltobacterales [45] from the phylum Eremiobacterota [46] (Fig. 3). Despite the presence of 71 Baltobacterales species-representative genomes in the GTDB R214 database (28th April 2023), all seven recovered genomes were identified as nMAGs. Many Eremiobacterota species were recently described by metagenome assembly from a number of boreal ecosystems such as Alaskan mosses [18], Canadian acidic springs [47], and Swedish peatlands [48]. Baltobacterales were highly prevalent in our samples, detected in 109/131 (∼83%) samples, and two *Nyctobacter* MAGs were of high quality (QS = 0.95 and 0.91 respectively; Supplementary Data 1). While there is little information on the ecological traits of Eremiobacterota, several genera within this group were shown to have the capacity for C fixation [39]. Our study supports this finding, as each Eremiobacterota we detected showed partial completeness in most prokaryotic carbon fixation pathway considered (Supplementary Fig. S2). Pessi *et al.* [50] detected *nif* homologues in the Eremiobacterota species Candidatus *Lamibacter sapmiensis*. However, we only found partial completeness of an assimilatory nitrate reduction module (M00531) in Candidatus *Velthaea versatilis*, whose nitrogen reduction capacity has already been well described [49]. Interestingly, all Eremiobacterota nMAGs except *Tumulicola* BrAA6 can encode either partial or complete pathways for anoxygenic photosynthesis, supporting findings by Ward *et al.* [45]. Additionally, we detected partial completeness of the methane oxidation pathway across all detected species within this phylum, supporting the findings of Holland-Moritz *et al.* [51], who identified only two cases of the methane oxidation marker gene *pmoA.* Moreover, Ray *et al*. [52] suggested Eremiobacterota might use atmospheric chemosynthesis by assimilating formaldehyde through the serine pathway. Our results support this hypothesis, as all Eremiobacterota we detected exhibited partial completeness of this pathway (see Supplementary Fig. S2). Our study thus contributes novel Eremiobacterota genomes of potentially undiscovered species and information about their metabolic potential, and further supports existing knowledge about moss-associated members of this clade.

### Bacterial species associated with moss species

In order to identify differentially abundant taxa across groups of samples, we used ANCOM-BC2 [53], which estimates log_2_-fold changes in absolute abundances between sample groups through pairwise comparisons. To assess differences in abundance across moss species, we used the Dunnett’s test, with *D. undulatum* as a reference group. *D. undulatum* was selected because of its contrasting bacterial community composition, which served as a baseline for evaluating differential abundant bacteria among moss species (Fig. 4a). A total of 28 species were found to be differentially abundant across the three *Polytrichum* species compared to *D. undulatum* (Fig. 4a; Supplementary Fig. S3). Of these 28 bacterial species, 24 were nMAGs, underlining the contribution of the assembly process to the depth of our analyses.

**Figure 4.**
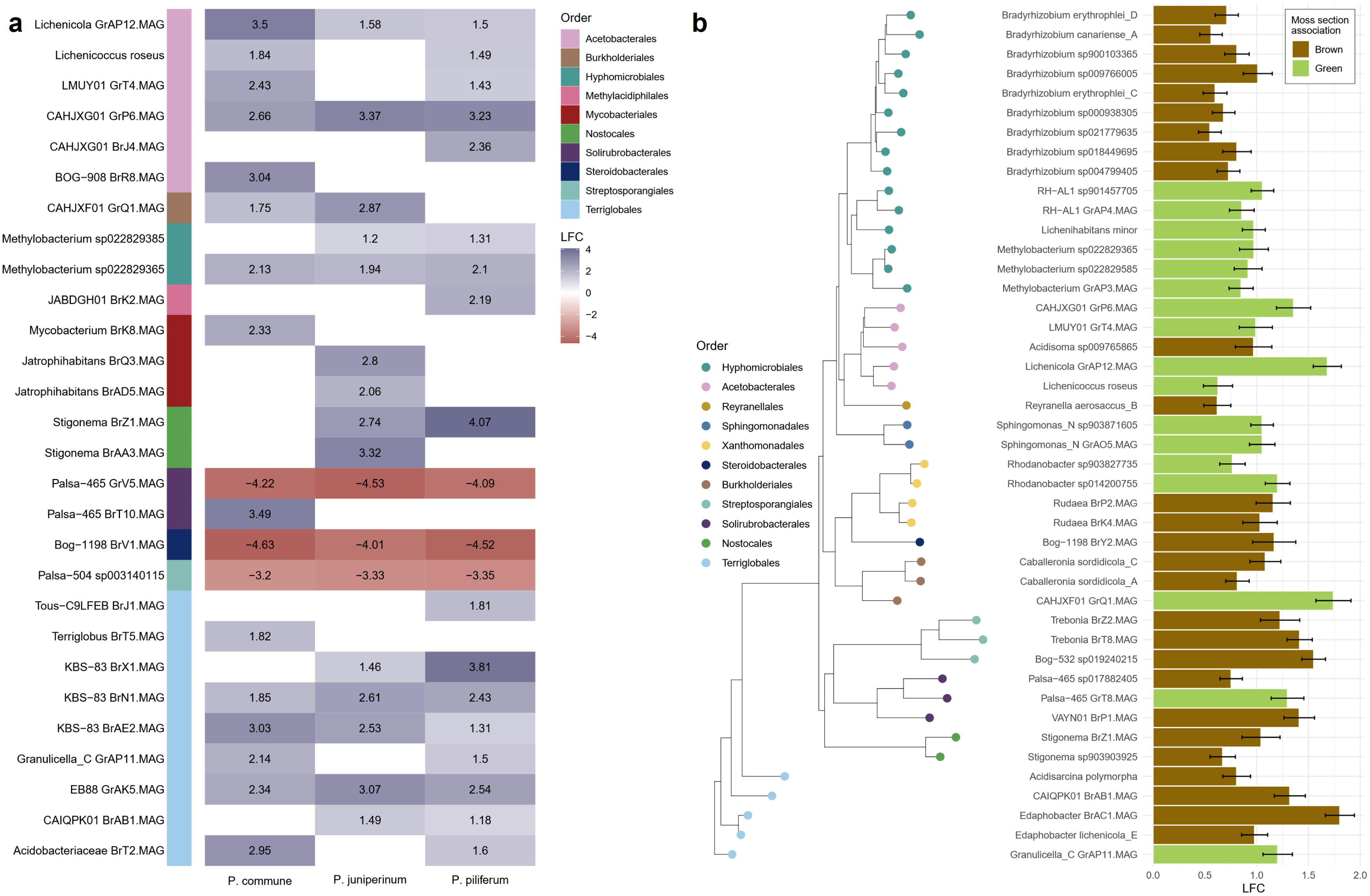
Differential abundance of bacterial species between moss species and between green and brown gametophyte sections. LFC: log_2_-fold change in absolute abundances, with all values showing significant differential abundance (p < 0.01). (a) Differentially abundant bacterial species in moss species relative to *D. undulatum*. LFC are relative to *D. undulatum*. Positive LFC (blue) indicates enrichment in specified moss species relative to *D. undulatum*, whereas negative LFC (red) indicates enrichment in *D. undulatum* relative to specified moss species. (b) Bacterial species differentially abundant in either green or brown sections of moss gametophyte. Bar colours indicate in which section (green or brown) the species was differentially enriched, with LFC calculated in relation to the reference group. Blank cells correspond to species for which no significant differences in abundance were detected, and therefore no LFC values reported.

We also performed a differential abundance test at the order level in a pairwise manner (i.e., without a reference host). Interestingly, we found that the order Baltobacterales was more abundant in *P. commune* by 2.7 to 3.3 log_2_-folds compared to all other moss species (Supplementary Fig. S3). Previous metagenomic studies showed that Baltobacterales encode genes necessary for hopanoid synthesis (a biochemical process in which hopanoid, steroid-like compounds, are produced) [49]. Hopanoid molecules were shown to contribute to bacterial tolerance to low pH soils [54], temperature stress [49], microbial defense [56], and are linked to N-fixation [57, 58]. Notably, *P. commune* is known for its antibacterial properties [60] and specimens from this moss species were sampled from acidic soils (Supplementary Data S2) in our study. The stress-tolerance properties of hopanoid structures in Baltobacterales could provide an adaptive advantage for colonization and persistence of bacteria in acidic and anti-microbial habitats as found in *P. commune* mosses.

Two *Stigonema* spp. (Nostocales) were differentially abundant in *P. juniperinum* and *P. piliferum* (relative to *D. undulatum*), but not in *P. commune* (Fig. 4a). One major difference between *Polytrichum* species is the habitat in which *P. commune* was growing in. We found that *P. commune* grew in a significantly more shaded environment than the other *Polytrichum* species (Supplementary Data S2), and *Stigonema* may be more sensitive to these differences in shading. Indeed, Jean *et al.* (2020) found that Nostocaceae abundance was negatively associated with canopy cover [8]. Furthermore, the aforementioned anti-microbial properties of *P. commune* could select against *Stigonema* spp. growth. Finally, *Stigonema*, known for having an alternative nitrogenase, is also common in boreal lichens (*Stereocaulon*) growing near *Polytrichum* and *D. undulatum*. [57]. The proximity of *Stigonema* to these moss species, combined with its unique functional adaptation, such as its alternative nitrogenase, may influence the abundance or composition of the bacterial communities associated with these mosses. In general, all Acetobacterales and Terriglobales species were positively associated with *Polytrichum* species (Fig. 4a). Of note, Terriglobales, one of the most overall relatively abundant species in our dataset (mean 20.8 ± 10.8% of attributed sequences across all samples, Fig. 2d), were also found to be highly present in sub-arctic and boreal peatland moss microbiomes [62].

Three species from the orders Solirubrobacterales, Steroidobacterales, and Streptosporangiales were found to be much more abundant in *D. undulatum* compared to *Polytrichum* hosts (3.20 to 4.63 log_2_-folds, Fig. 4a). The most closely related species from these orders were incidentally all species identified through metagenome assembly and were described in soil and roots in non-boreal ecosystems [52, 63]. A potential reason for the differences in bacterial communities between *Dicranum* and *Polytrichum* genera could be due to host environmental preferences (*D. undulatum* preferring moist soils), as well as moss N and phosphorus (P) content [64]. Although our results show that moss-associated bacterial communities are influenced by moss identity, experimental studies are needed to disentangle the simultaneous effects of biotic and abiotic conditions on moss-associated bacteria and on their contribution to ecosystem functions.

### Bacterial species associated with brown and green gametophyte sections

We identified 44 species as differentially abundant between green and brown gametophyte sections, of which 18 are nMAGs (Fig. 4b). These species are hereafter described as “associated” to the section in which they were more abundant. Our work identified 27 species as brown-associated, including all nine *Bradyrhizobia*, thus suggesting a strong specificity for brown gametophytes for members of this genus (Fig. 4b).

Bacterial species with the highest log_2_-fold change in brown gametophyte sections included three nMAGs from the genus *Edaphobacter*, uncultured species from the order Solirubrobacterales (*VAYN01*) and Terriglobales (*CAIQPK01*), as well as three uncultured species from the order Streptosporangiales (*Bog-532 sp019240215* and two *Trebonia* nMAGs; Fig. 4b). Various *Edaphobacter* species, such as *E. lichenicola sp. nov., Edaphobacter modestus gen. nov.*, and *Edaphobacter aggregans sp. nov.*, were previously described from alpine forest soils [65, 66]. Although Solirubrobacterales are poorly studied, researchers have detected members of this order in diverse ecosystems including agricultural lands [67] and Antarctic soils [68]. Several other brown-associated taxa, such as *Rudaea* spp., *CAIMXF01*, *Trebonia*, and *Acidisarcina*, are also mainly associated with soils [69–71]. Finally, two *Stigonema* species (cyanobacteria of the order Nostocales) were differentially abundant in brown gametophytes (Fig. 4b). In line with our findings, Renaudin *et al*. [10] found that cyanobacteria biomass increased towards the bottom of moss gametophyte, peaking at approximately 2-3 cm from the apex. In their study on the bacterial communities of two *Racomitrium* moss species, Klarenberg *et al.* [72] demonstrated that there is little transmission of bacteria from soil to moss gametophytes [69]. Meanwhile, Escolástico-Ortiz *et al.* [73] showed overlap between soil bacterial communities and the moss *Racomitrium lanuginosum*. However, the authors also revealed the presence of a core microbiome for moss gametophytes and soil, with each harbouring distinct, non-overlapping core bacterial communities. While we did not characterize the bacterial community of the surrounding soil, our finding of a number of soil species in the brown gametophyte section suggests this habitat may be colonized by both soil and moss-associated microbes.

In the green gametophyte section, nMAGs from the genera *CAHJXF01*, *CAHJXG01*, *Lichenicola*, and *Palsa-465* had the highest change in absolute abundance (log_2_-fold > 1.29; Fig. 4b) compared to the brown gametophyte section. Hay *et al*. [75] described MAGs of *CAHJXF01* and *CAHJXG01* from high Arctic glaciers, containing the gene form I RubisCo – an important gene for C fixation. We also found certain species of Hyphomicrobiales (Rhizobiales) to be green-associated, specifically from the genera *Methylobacteria*, *Lichenihabitans*, and *RH-AL1*, as well as all species of the genus *Bradyrhizobium* (Fig 4b). *Methylobacteria*, for example, were found in the core microbiome of boreal shrubs [76] and were shown to colonize non-boreal mosses (e.g., *Didymodon tectorum* and *Funaria hygrometrica* [77, 78]). Similarly, *Lichenihabitans minor* and *RH-AL1* spp. were shown to harbor genes associated with methanol consumption [74]. Although *Lichenihabitans minor* tends to be associated with lichens, the detection of this species in our samples is not surprising, given the prevalence of lichens in the habitat of boreal mosses [79]. *Lichenicola* and *Lichenicoccus* species, eponymously, can associate with lichens. For example, *Lichenicola cladoniae* was previously found in Antarctic lichens [80]. Overall, through our analysis, we observed the association of several species from genera such as *CAHJXF01*, *CAHJXG01*, *Methylobacteria*, *Lichenihabitans*, and *RH-AL1* spp. with the green gametophyte section. Future studies would be warranted to quantify their impact on C fixation and methanol consumption in boreal forests.

### Bacterial metabolic potential differs in green and brown gametophyte sections

Next, we investigated the metabolic potential of the gametophyte section-associated bacteria by analyzing which pathways their genomes have the potential to express. We focused on four pathway groups relevant to boreal ecosystem functioning: photosynthesis, nitrogen metabolism, methane metabolism, and carbon fixation.

First, we found that two brown-associated *Stigonema* species, also known as true-branched filamentous cyanobacteria, had partial pathway completeness for photosystems I and II (*Stigonema sp903903925* = 83.3% and *Stigonema* BrZ1.MAG = 66.7%) (Fig. 5; Supplementary Data S3). These were the only two differentially abundant species capable of at least partially encoding these pathways. Moreover, two *Stigonema* spp. stood out because of their partially complete N fixation (nifHDK *+* vnfDKGH) pathway (*Stigonema sp903903925* = 75% and *Stigonema* BrZ1.MAG = 50% completeness) (Fig. 5; Supplementary Data S3). This supports a previous finding in feathermosses, which showed *Stigonema* spp. to be important contributors to N fixation [81, 82]. *Stigonema* spp. also showed partial pathway completeness in all methane modules relevant to methanogenesis, and not methane oxidation. Although methanogenesis has not been shown to occur in any organisms except archaea [83], partial completeness of several methanogenesis pathways alludes to the potential for certain cyanobacteria to contribute to non-methanogenesis methane production (Fig. 5; Supplementary Data S3). Overall, our findings suggest that Nostocales species associated with the brown gametophyte section are photosynthetically active and have the potential for nitrogen fixation [85, 86].

**Figure 5.**
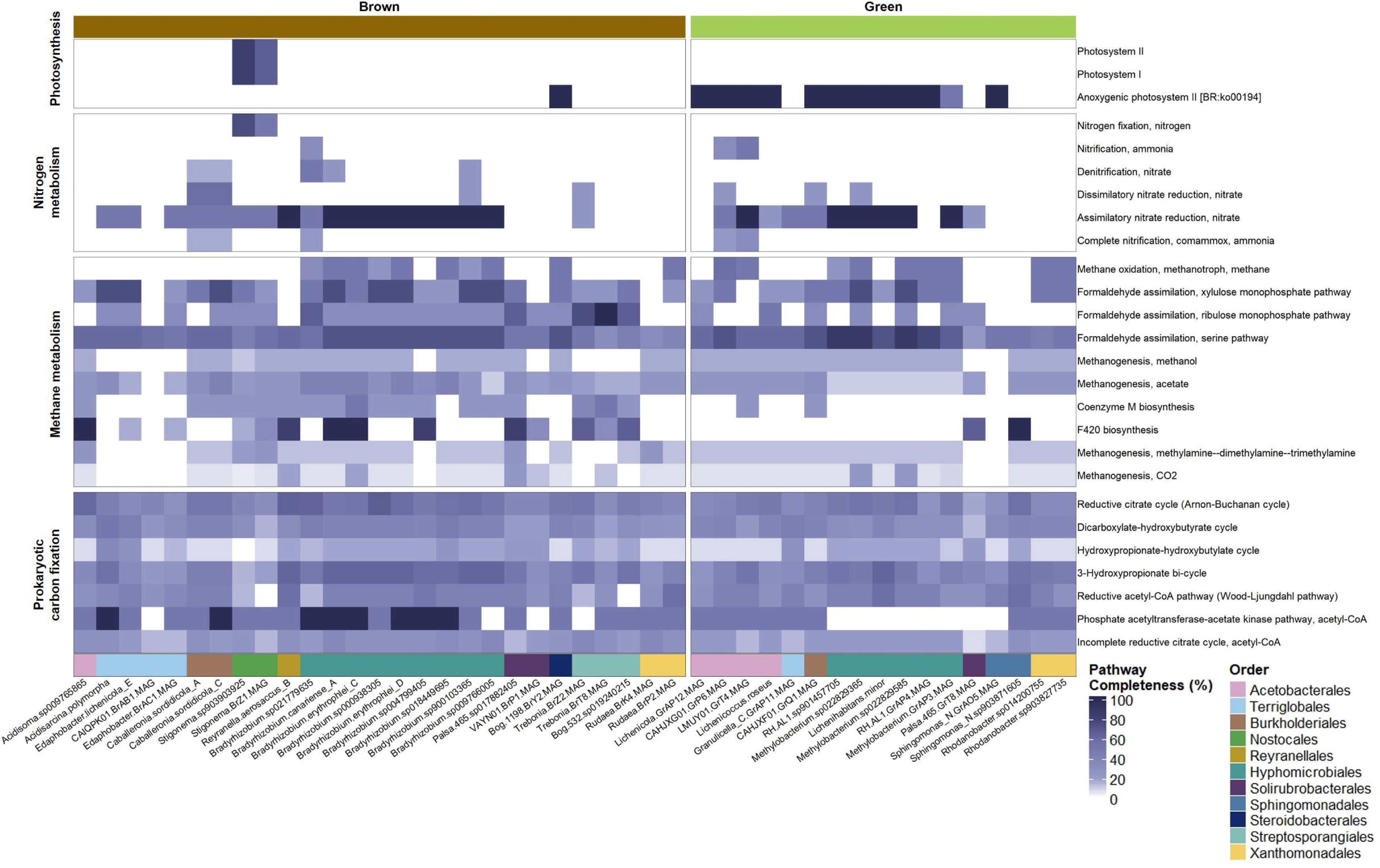
Metabolic abilities of the differentially abundant species in brown and green gametophyte sections. The pathway completeness of selected metabolic pathway groups, such as photosynthesis, nitrogen and methane metabolism, and prokaryotic carbon fixation, is indicated for each genome (blue scale). This figure also shows the taxonomic rank (e.g., bacterial order) for each of 44 differentially abundant genomes.

Second, we detected complete assimilatory nitrate reduction pathways in both green- and brown-associated Hyphomicrobiales (Rhizobiales), in line with earlier findings on nitrogen metabolic capacities of this clade (Fig. 5; Supplementary Data S3). Hyphomicrobiales taxa were previously shown to be N-fixers for nodule-producing leguminous plants [87] and Sphagnum mosses [17]. Interestingly, we found that nitrogen metabolism pathways were not significantly more complete in bacteria associated with either green or brown gametophyte sections. This suggests that the potential of the bacterial community to encode these pathways is not influenced by section type (Supplementary Table S1). Furthermore, most methane metabolic pathways were also partially complete in both green and brown-associated Hyphomicrobiales genomes (Fig. 5; Supplementary Data S3). For example, the methanogenesis acetate pathway was significantly more complete in brown-associated species of *Bradyrhizobium* (p-value = 0.021; Fig. 5; Supplementary Table S1). This aligns with the results of a previous study on *Sphagnum* mosses, which found that methanotrophy is primarily carried out by Hyphomicrobiales species [88]. Additionally, two of the three green-associated Methylobacteria showed potential to fully encode the anoxygenic photosystem II pathway, whereas the third one (one of the nMAGs) can only encodes 50% of the required genes (Fig. 5; Supplementary Data S3). Of note, the only photosynthesis pathway whose completeness was significantly associated with the green-associated species was also the anoxygenic photosystem II (p-value = 0.008 Supplementary Table S1). This is in line with recent findings that aerobic anoxygenic photosynthetic bacteria are present among plant phyllosphere microbial communities [89], including certain species of Methylobacteria [90]. Next, not a single green-associated Hyphomicrobiales encodes any gene of the phosphate acetyltransferase-acetate kinase pathway (carbon fixation), whereas among brown-associated Hyphomicrobiales, six *Bradyrhizobium* spp. encode 100% of the required genes (Fig. 5; Supplementary Data S3). In line with this finding, the phosphate acetyltransferase-acetate kinase pathway was also the only pathway significantly more complete in differentially brown-associated species (p-value = 0.012; Supplementary Table S1). This pathway is part of a central metabolism of bacteria [91]; its high completeness in *Bradyrhizobia* indicates the metabolic potential for C fixation [92]. To sum up, our findings suggest that green- and brown-associated Hyphomicrobiales provide a range of metabolic contributions such as C fixation, methane metabolism, and nitrate reduction to their ecosystem.

Third, brown-associated Burkholderiales (*Caballeronia* spp.) showed potential for denitrification activity, dissimilatory and assimilatory nitrate reduction, and a complete nitrification pathway (Fig. 5; Supplementary Data S3). However, the only green-associated Burkholderiales species (CAHJXF01 nMAG) lacked all three nitrification pathways (denitrification, nitrification, and complete nitrification, Fig. 5). While we did not find methane oxidation pathways in these species, they still showed completeness in several methane metabolism pathways, such as formaldehyde assimilation (Fig. 5; Supplementary Data S3). These findings suggest the distinct metabolic capabilities of Burkholderiales in different sections of the gametophyte [93, 94]. Curiously, pathways for coenzyme M (p-value = 0.015) and F420 biosynthesis (p-value = 0.169) were partially to fully complete in several brown-associated species (Fig. 5; Supplementary Table S1 and Supplementary Data S3). F420 is a costly co-enzyme to synthesize; nevertheless, Ney *et al*. [95] demonstrated it was commonly synthesized by aerobic bacteria. Meanwhile, coenzyme M is found in methanotrophic archaea and used by bacteria in metabolizing of different substrates, for example alkenes [96], which are present in boreal forest soils [97]. More interestingly, alkenes are known to be produced by bacteria associated with lichens [98]. In our sampling sites, lichens frequently grew alongside mosses, indicating that some of the bacterial taxa may be shared between the groups. Our results suggest that the presence of coenzyme M in bacteria associated with brown gametophyte sections may be attributed to their utilization of alkenes commonly found in lichens.

Overall, our findings demonstrate the diverse metabolic potential and adaptations of bacterial species associated with different sections of moss gametophyte, shedding light on their roles in N cycling, methane metabolism and C fixation in boreal forests.

### Conclusion

In this study, using a whole metagenome sequencing approach, we demonstrated that moss genus, moss species, gametophyte section, and, to a lesser extent, soil pH and soil temperature are significant drivers of bacterial community composition and diversity (Tables 1-2; Fig. 2). Through metagenomic assembly, we identified 110 potentially novel bacterial species (Fig. 3) and characterized distinct bacterial communities associated with four different moss species (Fig. 4). We found that bacterial communities in green and brown gametophyte sections occupy unique functional niches (Fig. 5), with partial partial overlap. Our results show that the brown gametophyte section, despite their senescent nature, can support bacterial communities that contribute to ecological functions such as nitrogen fixation, carbon fixation, and methane metabolism, while green-associated bacteria exhibit potential for anoxygenic photosynthesis.

To build on these findings, several key areas for future research emerge. First, validating the specific metabolic pathways of the 110 potentially novel bacterial species and their contributions to processes like nitrogen fixation and carbon sequestration is important. For example, experimental studies and transcriptomic studies could help to identify which genes are actively transcribed under different environmental conditions. Second, exploring the interactions between moss-associated bacteria, fungi, and other microbial members could provide more insights into their joint roles in ecosystem functioning. Gaining a clear understanding of moss metabolites could further elucidate the potential co-evolutionary or symbiotic relationships between non-cyanobacterial microbial communities and mosses, revealing how metabolites influence microbial interactions and ecosystem processes. To summarize, our results contribute to our understanding of how moss-bacteria interactions contribute to ecosystem functions, such as nutrient cycling and carbon sequestration, emphasizing the importance of both green and brown gametophyte section-associated bacteria in these processes.

## Methods

### Study area

This study was conducted within the Eeyou Istchee (James Bay) region of Northern Québec (Fig. 1a; N51.7040721°, W76.0521982°). The sampling area is part of Québec’s boreal forest biome and is dominated by closed-crown forest stands of Jack pine (*Pinus banksiana*) and black spruce (*Picea mariana*) [99]. The monthly average annual temperature of this region is 0.3LJ, while mean annual precipitation is 741 mm [100]. The soils are predominantly acidic and nutrient poor [101].

### Study design and moss species

Samples were taken from four acrocarpous (stems harbor archegonia at the top) moss species: *Polytrichum juniperinum*, *Polytrichum commune*, *Polytrichum piliferum*, and *Dicranum undulatum* (Fig. 1b), which are native to the Eeyou-Istchee region of Québec [28]. These were sampled in September 2021, at five different locations (Fig. 1a). Three microsites, each 5 to 10 meters apart, were randomly selected and then, at each microsite, 3-by-3-inch moss patches were sampled directly (Fig. 1a). At each microsite/species, we measured soil temperature, by inserting the probe of the Fisherbrand^TM^ Envirometer^TM^ directly in the soil of where the moss was sampled after it had been removed. Soil pH was measured in the laboratory using the method described by Hendershot, Lalande, and Duquette [102]. Mosses were first visually identified to species level in the field and then confirmed by microscopy. For each moss species, samples were taken from both the brown and green sections of the gametophyte, which were determined visually (Fig. 1c). To maintain the integrity of the samples, the green and brown sections were separated while on site, using ethanol and DNA-away treated tweezers and scissors. Samples were placed in sterile 50 mL tubes and immediately stored at -20LJ for the return trip to the laboratory, where they were stored at - 80LJ. A total of 131 samples were collected for brown and green gametophyte sections across *P. juniperinum* (N_brown=_ 24, N_green_ = 22), *P. commune* (N_brown=_ 9, N_green_ = 9), *P. piliferum* (N_brown=_ 26, N_green_ = 26), and *D. undulatum* (N_brown=_ 8, N_green_= 7) from five sites (see Supplementary Table S2 for sample breakdown).

### DNA extraction and whole metagenome sequencing (WMS)

Green and brown gametophyte sections were processed separately, but in the same manner. DNA extraction of moss samples was performed as modified versions of the protocols used by Holland-Moritz *et al.* [4] and Klarenberg *et al.* [16]. Using tweezers and scissors sterilized with 80% ethanol solution and ‘DNA Away’ solution (Thermo Fisher Scientific, MA, USA), 0.25 g of each moss sample were weighed out and divided into two sterile 2 mL bead beating tubes that also contained three 2.3 mM BioSpec sterilized stainless-steel beads. These tubes were then frozen in liquid N_2_ for approximately 30s and mosses were broken down using the FisherbrandTM Bead Mill 24 Homogenizer for 30s at 1.75 m/s. DNA was subsequently extracted using the QIAGEN DNEasy PowerSoil DNA Extraction Kit with some minor changes. Each moss sample was divided into two 2 mL PowerBead extraction tubes. The kit buffer and beads were added to each tube, along with 60 uL of C1 Solution from the kit. All tubes with buffer were homogenized at 3.25 m/s for 30s and then at 2.40 m/s for 10 minutes using the FisherbrandTM Bead Mill 24 Homogenizer. After centrifugation as per the extraction kit protocol, supernatant from both tubes of one sample were combined in one tube with 250 uL of C2 Solution from the extraction kit. The rest of the extraction followed the manufacturer’s instructions. All extractions were performed under a Type A2 Bio Safety Cabinet. Extracted DNA were subsequently quantified using the NanoDropTM One Spectrophotometer (Thermo Fisher Scientific, MA, USA). Samples under 20 ng/uL or with A260/230 ratios less than 1.8 were tested for presence of bacteria by PCR amplification of the V4-V5 region of bacterial 16S rRNA using 515F (GTGCCAGCMGCCGCGGTAA) as forward primer and 806R (GGACTACHVGGGTWTCTAAT) as reverse primer. The 25 uL PCR reaction mix for these tests consisted of 5 uL of 1X GC Buffer, 0.5 uL of 10uM dNTPs, 0.5 uL each of 10uM forward and reverse primers, 0.25 uL HotStart DNA polymerase and 17.25 uL RNAse free MilliQ water. PCR thermocycling conditions were an initial denaturation at 94LJ for 3 minutes, followed by 35 cycles of: denaturation at 95LJ for 30s, annealing at 50LJ for 30s, elongation at 72LJ for 1-minute 30s; lastly there was a final elongation step at 72LJ for 10 minutes. All tested samples showed amplification at the predicted length of 290 base pairs.

In total, 131 metagenomic libraries were generated. Samples were sequenced on a single lane of Illumina NovaSeq6000 with 2 × 150bp chemistry at Génome Québec (Montréal, Canada).

### Bioinformatic analyses

Samples preprocessing (quality control), assembly, MAG generation and annotation, as well as novelty analyses, were performed on high-performance computer with the aid of SLURM workload management for parallel processing. The pipeline for these steps is written in bash. In the next steps, the R programming language was used for diversity and differential abundance analyses, as well as production of figures and tables. R packages are listed in Supplementary Table S4, along with all bioinformatic software (version and non-default options, when applicable) outlined in the following methods. The complete workflow is provided in Supplementary Fig. S4.

### Sample quality control

The raw sample library size was 14.8 × 10^6^ ± 2.2 × 10^6^ reads. Raw reads were trimmed and quality-controlled using Kneaddata 0.12.0, keeping reads with a minimum length of 50 bp and an average quality score of 30 over a 4-nucleotide sliding window. At the time of the experiment, all four moss species sampled in this study lacked reference genomes. Thus, host DNA was partially removed by aligning reads against the closely related genome of *Physcomitrella patens*, as well as the human genome GRCh38 to remove human contamination, using Bowtie 2.4.4 (from within the Kneaddata pipeline). On average, clean samples contained 10 × 10^6^ ± 1.6 × 10^6^ paired-end reads.

### Choice of a co-assembly strategy

Metagenome assembly quality depends on depth of coverage of each genome within samples [103]. A common strategy is to co-assemble samples based on biological similarities, by pooling sequences submitted to the assembler, increasing the coverage of genomes, and therefore retrieving more good quality genomes [104]. In this study, we pooled samples (i.e., concatenated the *.fastq* metagenome files) taken from the same moss species, gametophyte section, and microsite into 46 sample pools. We applied the pipeline described in the following section identically to both assembly and co-assembly strategies. Co-assembly generated 157 MAGs with completeness > 50% and contamination < 10%, compared to 139 through regular sample-level assembly. Given this, we focused on the MAGs generated by co-assembly, which had a slightly higher average completeness, increasing from 73.6% ± 16.5 to 74.2% ± 15.4%.

### Metagenome co-assembly and MAG generation

Samples were co-assembled using MetaWRAP modules [105]. First, paired-end sequences were assembled using metaSPAdes 3.15.4 and re-assembled unassembled reads using MEGAHIT 1.2.9. Contigs were then binned with three binners (MetaBAT 2.12.1, CONCOCT 1.1.0, and MaxBin 2.2.4). Afterwards, these three sets of bins were hybridized with the MetaWRAP 1.3 *bin_refinement* module to produce a final set of refined bins for each co-assembly. Bin quality was then assessed using CheckM 1.0.18 and bins with completeness < 50% and contamination > 10% were excluded [106], yielding a set of 260 bins (5.7 ± 3.3 bins per co-assembly).

To obtain a unique set of species-level MAGs for the whole study, bins were pooled from all co-assemblies and dereplicated them using dRep 3.4.0 [107]. dRep uses a two-step clustering strategy, whereby bins are first clustered using Mash 2.3 [108]; then, within each cluster, bins are dereplicated at a 95% average nucleotide identity (ANI) using MUMmer 3.23 [109]. GUNC 1.0.5 [110] was used to identify chimeric MAGs, which were then removed from the final set of species-level metagenome-assembled genomes (MAGs).

### MAG novelty analysis

MAGs which potentially belonged to species with no published reference genome, and therefore were absent from the reference database, were identified by applying three bioinformatic validations. First, we taxonomically annotated MAGs using GTDB-Tk [111]. Second, we calculated the Mash distance between these 156 MAGs and the 31,332 non-redundant bacterial and archaeal genomes from EnsemblBacteria. Third, since Mash distance could be underestimated when comparing incomplete MAGs, we used skani 0.2.1 [112] to compute ANI between these MAGs and the 85,205 GTDB species representative genomes. MAGs were labelled as novel MAGs (nMAGs) if they had an ANI < 0.95 having a Mash distance > 0.05 and no species-level taxonomy assignment. Additionally, after adding the nMAGs to the reference genome index as described below, two were dropped because they were not detected by Sourmash 4.7.0 [113]. nMAGs were only kept if they exhibited an assembly quality score (defined as Completeness – 5 × Contamination) > 0.50, yielding a final set of 110 nMAGs.

### nMAG annotations

nMAGs were further annotated by predicting their coding sequences using Prodigal 2.6.3 [114] and passing these to the MicrobeAnnotator 2.0.5 [115] pipeline, generating an estimation of their metabolic potential through pathway completeness reconstruction (see *Pathway completeness analysis* below for further details). Finally, phylogeny was estimated using PhyloPhlAn 3.0.3 [116] and phylogenetic trees built using ggtree 3.10.0 [117] and ggtreeExtra 1.12.0, allowing a cross-validation with their attributed GTDB taxonomy. nMAGs statistics (length, L50, number of contigs and GC contents) were assessed using bbmap 38.86.

### Taxonomy characterization and abundance estimation of bacterial community

Sample community composition and relative abundance at the species level was estimated using Sourmash gather [113] with a *k*-mer size of 31, using the GTDB representative species genome index [111] to which nMAGs were added. This approach also measures a sample’s containment, which represents the proportion of sample *k*-mers found in the smallest set of reference genomes required to cover all sample *k*-mers with an exact match. Of note, *k*-mer-based community profiling tools generate a *sequence* abundance table; these abundance values are not normalized for genome length (as opposed to marker-gene-based tools), making them distinct from taxonomic abundance [118]. However, the necessity to introduce nMAGs in the reference database precludes the use of marker-gene referencing for taxonomic assignment.

Containment was used to estimate the improvement in sample coverage caused by the addition of nMAGs to the GTDB R214 reference index. The GenBank 2022-03 reference index (comprising 1.22 M microbial genomes) was also compared, and the containment increase is reported for both scenarios (Supplementary Fig. S1). Finally, the sequence abundance matrix was built into a phyloseq (version 1.46.0) object using the raw counts; this object was then used as a starting point for all subsequent analyses.

### Diversity analyses

All statistical analyses and plots were made using the open-access software R 4.2.1 [119]. For community composition (multivariate) analyses, uneven sequencing depth and heteroskedasticity were addressed by applying variance stabilizing transformations (VST, DESeq2 1.42 [120]) to the abundance matrix. This matrix was then used to estimate pairwise Bray-Curtis dissimilarity between samples [121], and to compute principal coordinates using the *capscale* function in the vegan package (version 2.6-4) [122]. Finally, the *adonis2* function from the vegan package [122] was used to run PERMANOVAs, assessing the sensitivity of model outputs to the order of its parameters.

In brief, we explored the effect of moss genus, moss species, gametophyte section, and soil abiotic factors on bacterial community composition. We ran models with and without *Dicranum* samples, thus testing the differences in bacterial community composition between moss genera and among *Polytrichum* spp. (*P. juniperinum*, *P. commune*, and *P. piliferum*), respectively. This approach allowed us to disentangle whether closely related moss species are likely to harbor more similar microbial taxa than more distantly related ones, because of shared physiological traits and similar ecological niches. In the models with *Dicranum* samples, we modelled multivariate bacterial community composition as a function of categorical variables ‘*moss genus*’, ‘*moss species*’ nested within ‘*moss genus*’, ‘*gametophyte section*’, and continuous variables ‘*soil pH*’ and ‘*soil temperature*’, and their two-way interactions. We nested ‘*moss species’* within the genera to account for the hierarchical taxonomic structure and to explore species-specific effects on bacterial community composition and alpha diversity (see below). In the models without *Dicranum* samples, we modelled multivariate bacterial community composition as a function of categorical variables ‘*moss species*’, ‘*gametophyte section*’, and continuous variables ‘*soil pH*’ and ‘*soil temperature*’, and their two-way interactions. A fixed effect ‘*moss species*’ was included in the models, to quantify the portion of variance in bacterial community composition explained by *Polytrichum* spp. In all multivariate models, permutations were restricted within blocks (i.e., ‘*Location’*), using the function *how* of the permute package [123].

For alpha diversity analyses, reads were rarefied to 24,900 (slightly less than the lowest sample sum of reads) using *rarefy_even_depth* (phyloseq) to account for uneven sequencing depth [40, 124]. Linear models were run on Shannon index using function *lmer* in the lme4 1.1-35.1 [125] package, then the *anova* function in the car 3.1-2 package was used to assess model significance. Final models were further simplified by removing insignificant interactions. For each of the alpha diversity models, the functions *simulateResiduals* and *plot* from the DHARMa 0.4.6 package were used to assess model fit [126].

To explore the differences in bacterial diversity among moss identities, gametophyte section and abiotic factors, bacterial diversity (Shannon index) was modelled as a function of categorical variables ‘moss *genus*’, ‘*moss species*’ (nested within ‘*moss genus*’), ‘*gametophyte section*’, and continuous variables ‘*soil pH*’ and ‘*soil temperature*’, and their two-way interactions. Finally, models excluding *Dicranum* samples were run to confirm effect persistence intra-genera, for example by including fixed factor ‘*moss species*’. In all models, to account for variation among sampling locations, the random effect ‘*Location’* was included, and to account for nested design, we nested ‘*Microsite’* under ‘*Location’*. See Supplementary Table S3 for more details on model structures, fixed and random effects tested, and link functions used.

### Differential abundance analyses (DAA)

To identify species associated with moss species and gametophyte section, DAA were performed using ANCOMBC 2.4.0 package [53]. This approach was chosen because it takes into account the compositionality of next-generation sequencing data (i.e., sequencing data is commonly expressed in terms of relative abundances, meaning they sum up to an arbitrary constant and reside in a simplex mathematical space) [127], controls for false discovery rate by accounting for multiple testing and multiple group comparisons, and corrects for both taxon- and sample-specific biases (i.e., differences in extraction efficiency across taxa and sampling fraction across samples). In brief, differentially abundant taxa were identified between (1) gametophyte section (green vs. brown), controlling for host moss species, soil pH, and soil temperature; and (2) host moss species, controlling for gametophyte section, using Dunnett’s test with *D. undulatum* as a reference group. In the latter, environmental covariates were omitted because they were correlated with location (SoilTemp ∼ Location, R^2^ = 0.58, p < 1e-10) and moss species (SoilpH ∼ Host, R^2^ = 0.62, p-value < 1e-10), and location is significantly associated with moss species (χ^2^ = 179.1, p-value < 1e-10). For both approaches, we only considered species present in at least 10% (n = 13) sample [128]. The Holm procedure was used to account for multiple comparisons. We only reported results that passed ANCOMBC’s sensitivity test to pseudo-count addition and had an adjusted p-value < 0.01.

### Pathway completeness analysis

Pathway completeness is a value from 0 to 100% representing the proportion of proteins making up a given pathway (or module) whose coding sequence could be found in a genome. The relationship between pathway completeness and gametophyte green and brown sections was investigated for the 44 differentially abundant species. The tests were performed on pathways from four groups of interest: nitrogen metabolism, photosynthesis, methane metabolism, and carbon fixation (Supplementary Table S1), as described in Kyoto Encyclopedia of Genes and Genomes (KEGG) [129]. Pathways for which completeness was ≤ 20% in more than 80% of the 44 DA species considered were excluded, yielding a subset of 23 pathways. Since pathway completeness correlates with genome completeness [130], we controlled for genome completeness (attributing 100% to non-MAGs). Specifically, the model *Module proteins (%) ∼ Gametophyte sections (green or brown) + MAG completeness* was tested, using a generalized linear model with a quasibinomial distribution and a logit link function to fit the response variable, which is a proportion that can include both boundaries (0% and 100%). P-values were corrected using the Holm method and only pathways with adjusted p-values < 0.05 were reported as significant. The heatmap in Fig. 5 was generated using the ComplexHeatmap 2.18.0package and shows a slightly different set of pathways compared to Supplementary Table S1, as certain pathways that were excluded for the statistical test on the basis of completeness-prevalence (as mentioned above) were still of interest for the discussion.

### Environmental DNA contamination assessment

The quasi absence of non-bacterial genomes in our assemblies prompted us to investigate the high-level taxonomy of sample reads. 100,000 reads per sample were randomly selected and used to perform a BLASTn search against the entire *ncbi_nt* database (across all Kingdoms) using BLAST+ 2.12.0. Although this approach cannot be used to estimate the relative abundance of non-microbial reads, our data showed that numerous reads belonged to eukaryotic, non-microbial species (see Supplementary Fig. S5-6). Therefore, the effective sequencing depth of bacterial species, even after adding our nMAGs, is lower than the sample sequencing depth. Of note, as the focus of this work being on bacteria, we designed the metagenome assembly pipeline to focus on metagenomic bins of bacterial origin, implicitly excluding eukaryotes that might have been assembled.

Furthermore, since no reference genome for the studied mosses was available at the time the assembly and analyses were performed, we assessed *a posteriori* the potential of host DNA to have contaminated our MAGs. To this end, reads from pooled sample were mapped to every MAG separately using Bowtie2 *(--very-sensitive*). Reads aligning with MAGs were then mapped to the recently published genome of *P. commune* (GCA_950295325.1). With one exception, the proportion of sample reads mapping to both, any MAG and *P. commune*’s genome, was extremely small, at 0.02 ± 0.07%. The exceptional MAG, *Green_N.bin.3*, had 3.7% of reads mapping to it also mapping to *P. commune*’s genome. This MAG was annotated as a *Buchnera* species (Supplementary Data 1); a phylogenetically close species, *Buchnera aphidicola*, has a particularly small genome (618 kbp), which could explain why the proportion of its genome conserved with a plant could be higher than for a bacteria with a more common genome size. Nevertheless, we tested our differential abundance analyses anew by excluding this MAG from the dataset, and the results presented in this work remained completely unaltered, as this MAG had not been found to be differentially abundant across gametophyte section or host mosses.

## Acknowledgements

First and foremost, the authors would like to thank the Cree Community of Nemaska for welcoming our team and allowing us to sample on their land. We thank Nemaska Lithium for their collaboration and support of the project. We would also like to acknowledge J. Chamard and members of S. Roy’s Lab (L. Garneau, J. Beaudin, O. Alix-Paré, and S. Malo) for the much-appreciated help with sampling and moss dissections, E. Lussier for extensive support with laboratory work, J.-F. Lucier for assistance with metagenomic analysis, and M. Desrochers for map generation. We are grateful to Génome Québec and to the Digital Research Alliance of Canada for providing us with access to super computers to perform our bioinformatic analyses. We also thank J. Beaudin, P. Moffett, and P. B. Beauregard for critical reading of the manuscript. We are grateful to Isabel Ramirez for the beautiful depictions of mosses presented in Fig. 1b. This work was supported by funding from (1) Fonds de recherche du Québec – Nature et technologies (FRQNT) Programme partenariat sur le développement durable du secteur minier (Grant 285683 lead PI Pr. Roy and co-PI Pr. Laforest-Lapointe); and (2) Pr. Laforest-Lapointe’s Canada Research Chair on Applied Microbial Ecology.

## Data availability statement

Scripts for all bioinformatic and statistical analyses, along with abundance tables and metadata used for analysis, were made public in an online repository (https://github.com/jorondo1/borealMoss). Raw sequencing reads, assemblies, and MAGs were deposited in the European Nucleotide Archive (ENA) at EMBL-EBI under accession number PRJEB76464.

## Author contributions

SI, SR, and ILL designed the experiment. SI conducted the sample collection, processing, and molecular work. JRL developed the bioinformatic workflow and conducted the bioinformatic analyses with the contribution from SI, MF, and ILL. SI and JRL analyzed the data with support from MF and ILL. SI wrote the first draft with the contribution from JRL, MF, ILL, and SR. All authors reviewed the final manuscript.

## Additional Information

Authors declare no competing interests.

