## Supplementary Tables and Figures for "Boreal moss-microbe interactions are revealed through metagenome assembly of novel bacterial species"

The following supporting documents are available for this paper:

**Supplementary Data:**

**Supplement Data 1.** Table shows 110 novel MAGs described in the study. Class and Genus = taxonomic classification; L50 = smallest number of contigs whose length sum makes up half of genome size; N50 = the sequence length of the shortest contig at 50% of the total assembly length; № of contigs = number of contigs in each MAG. Quality control identifiers: GC = guanine-cytosine (GC) content of a genome sequence; MBP = assembly span/genome size; comp = genome completness; cont = genome contamination; QS (aka Phred score) = measure for base quality.

**Supplementary Data 2.** Metadata for mosses and locations. Table shows sampling time, gametophyte sections, sampling locations and moss genus and species. It also provides data for measured abiotic factors such as soil pH and temperature.

**Supplementary Data 3**. Table shows pathway completeness (%) for each of 44 differentially abundant species and pathway.

**Supplementary Data 4.** Table shows pathway completeness (%) for each Eremiobacteria MAG from our total dataset.

**Supplementary Table S1**. Relationship between pathway completeness (percentage of module proteins identified in genomes) and gametophyte sections (green and brown) among 44 differentially abundant species. Analysis included 22 pathways retained after filtering those with ≤ 20% completeness in more than 80% of genomes. Completeness was assessed using a generalized linear model with a quasibinomial distribution, controlling for MAG completeness. Shown are pathways with significant associations (p_adj_ < 0.05) after Holm correction. In the statistical test, the reference compartment-association category is brown-associated bacteria. Therefore, positive estimates mean the pathway is significantly more complete in green gametophyte section, while negative estimates indicate it is more complete in the brown gametophyte section.

| Pathway ID | Pathway name | Pathway group | Estimate | *p*-value (adj) |  |
| --- | --- | --- | --- | --- | --- |
| M00165 | Reductive pentose phosphate cycle (Calvin cycle) | Carbon fixation | -0.297 | 0.301 |  |
| M00166 | Reductive pentose phosphate cycle, ribulose-5P => glyceraldehyde-3P | Carbon fixation | -0.317 | 0.347 |  |
| M00167 | Reductive pentose phosphate cycle, glyceraldehyde-3P => ribulose-5P | Carbon fixation | -0.352 | 0.442 |  |
| M00168 | CAM (Crassulacean acid metabolism), dark | Carbon fixation | 0.215 | 0.828 |  |
| M00169 | CAM (Crassulacean acid metabolism), light | Carbon fixation | -1.403 | 0.069 |  |
| M00170 | C4-dicarboxylic acid cycle, phosphoenolpyruvate carboxykinase type | Carbon fixation | 0.190 | 0.515 |  |
| M00172 | C4-dicarboxylic acid cycle, NADP - malic enzyme type | Carbon fixation | -0.380 | 0.231 |  |
| M00173 | Reductive citrate cycle (Arnon-Buchanan cycle) | Carbon fixation | -0.265 | 0.071 |  |
| M00374 | Dicarboxylate-hydroxybutyrate cycle | Carbon fixation | -0.061 | 0.611 |  |
| M00376 | 3-Hydroxypropionate bi-cycle | Carbon fixation | -0.112 | 0.543 |  |
| M00377 | Reductive acetyl-CoA pathway (Wood-Ljungdahl pathway) | Carbon fixation | 0.216 | 0.301 |  |
| M00579 | Phosphate acetyltransferase-acetate kinase pathway, acetyl-CoA => acetate | Carbon fixation | -1.371 | 0.012 | * |
| M00620 | Incomplete reductive citrate cycle, acetyl-CoA => oxoglutarate | Carbon fixation | -0.357 | 0.008 | ** |
| M00174 | Methane oxidation, methanotroph, methane => formaldehyde | Methane metabolism | 0.612 | 0.299 |  |
| M00344 | Formaldehyde assimilation, xylulose monophosphate pathway | Methane metabolism | -0.288 | 0.473 |  |
| M00345 | Formaldehyde assimilation, ribulose monophosphate pathway | Methane metabolism | -1.090 | 0.039 | * |
| M00346 | Formaldehyde assimilation, serine pathway | Methane metabolism | 0.241 | 0.299 |  |
| M00357 | Methanogenesis, acetate => methane | Methane metabolism | -0.477 | 0.021 | * |
| M00358 | Coenzyme M biosynthesis | Methane metabolism | -2.046 | 0.015 | * |
| M00378 | F420 biosynthesis | Methane metabolism | -1.412 | 0.169 |  |
| M00531 | Assimilatory nitrate reduction, nitrate => ammonia | Nitrogen metabolism | -0.050 | 0.928 |  |
| M00597 | Anoxygenic photosystem II [BR:ko00194] | Photosynthesis | 4.000 | 0.008 | ** |
| M00165 | Reductive pentose phosphate cycle (Calvin cycle) | Carbon fixation | -0.297 | 0.301 |  |

|  | Chemin PK  (51.56852°N; 75.54803°W) | Eastmain S1  (51.72042°N; 76.03975°W) | Eastmain S2  (51.72956°N; 76.01588°W) | Nemaska Inter.  (51.68096°N; 76.15626°W) | Pine Forest  (51.68686°N; 76.11511°W) |
| --- | --- | --- | --- | --- | --- |
| Moss species |  |  |  |  |  |
| *P. juniperinum* |  | n = 6 |  | n = 9 | n = 9 |
| *P. piliferum* | n = 9 | n = 9 |  |  | n = 9 |
| *P. commune* |  |  | n = 9 |  |  |
| *D. undulatum* |  |  | n = 9 |  |  |

Supplementary Table S2. Distribution of moss sample counts across five locations.

| Model type | Diversity | Data | Model predictors | Normalization or transformations | Responses examined | Model used |
| --- | --- | --- | --- | --- | --- | --- |
| Full models | *Alpha* | *P. juniperinun, P. commune, P. piliferum*, *&* *D. undulatum* collected from five sites in Eeyou Istchee (n=131, see Table S1 for a breakdown of the number of individuals collected) | Genus + Genus/Species + Section + Soil Temp. +  Section × Soil pH +  Genus × Section +  Species × Section +  Genus × Soil Temp. +  Species × Soil Temp. + Section × Soil Temperature + (1\|Microsite/Location) | Rarefied to 24900 sequences | Shannon diversity | Linear mixed effects model  (lmer) |
|  | *Beta* |  | Genus + Genus/Species + Section + Soil Temp. + Soil pH + Genus × Section + Species × Section +  Genus × Soil Temp. + Species × Soil Temp. +  Genus × Soil pH + Species × SoilpH +  Section × Soil pH + Section × Soil Temp.  Soil Temp. × Soil pH  [Restricted permutations within block ‘Location’] | Variance stabilizing transformed (VST) | Bray-Curtis dissimilarity | PERMANOVA  (adonis2) |
| Models without *Dicranum* | *Alpha* | *P. juniperinun, P. commune, P. piliferum*, collected from five sites in Eeyou Istchee (n=121, see Table S1 for a breakdown of the number of individuals collected) | Species + Section + Soil Temp. +  Section × Soil Temperature +  Section × Soil pH + Species × Section +  Species × Soil Temperature +  (1\|Microsite/Location) | Rarefied to 24900 sequences | Shannon diversity | Linear mixed effects model  (lmer) |
|  | *Beta* |  | Species + Section + Soil Temp. + Soil pH +  Species × Section + Species × Soil Temp. +  Species × Soil pH +  Section × SoilTemp + Section × Soil pH +  Soil Temp. × Soil pH  [Restricted permutations within block ‘Location’] | Variance stabilizing transformed (VST) | Bray-Curtis dissimilarity | PERMANOVA  (adonis2) |

Supplementary Table S3. Summary of model structures, responses examined, model predictors and normalization approaches used in alpha- and beta-diversity analyses.

Supplementary Table S4. Summary of bioinformatics steps, tools, versions, reference databases and non-default options used for metagenome co-assembly and MAG generation.

| Bioinformatics steps | Tool | Version | Reference database | | Non-default parameters |
| --- | --- | --- | --- | --- | --- |
| Preprocessing | Kneaddata | 0.12.0 | n/a | | Trimmomatic options: SLIDINGWINDOW:4:30 MINLEN:50 |
|  | bowtie2 | 2.4.4 |  | | --very-sensitive-local |
| Assembly | metaSPAdes | 3.15.4 | n/a | | k: [21, 33, 55], Mode: ONLY assembling (without read error correction) |
|  | MEGAHIT | 1.2.9 | n/a | |  |
| Binning | metaBAT2 | 2.12.1 | n/a | | minContig 1500, minCV 1.0, minCVSum 1.0, maxP 95%, minS 60, and maxEdges 200 |
|  | MaxBin | 2.2.4 | n/a | | Minimum contig length 1000 |
|  | CONCOCT | 1.1.0 | n/a | |  |
| Bin refinement | CheckM | 1.0.18 | n/a | | >50% completion and <10% contamination |
|  | metaWRAP | 1.3 | n/a | |  |
| Dereplication | dRep | 3.4.0 | n/a | | --P_ani 0.90 --S_ani 0.95 --cov_thresh 0.1 --completeness 0.5 --contamination 0.10 --S_algo ANImf |
| Chimera removal | GUNC | 1.0.5 | gunc_db_progenomes2.1 | | considered a chimera when "clade_separation_score > 0.45 && contamination_portion > 0.05 && reference_representation_score > 0.5" |
| Novelty analysis | Mash | 2.3 | genbank_57_bacteria | | dist -d 0.05 |
|  | skani | 0.2.1 | GTDB R214 | |  |
| MAG statistics | bbmap | 38.86 | n/a | |  |
| Metabolic annotation | MicrobeAnnotator | 2.0.5 | RefSeq, Swissprot, Trembl | | --refine |
|  | Prodigal | 2.6.3 |  | |  |
| Taxonomic profiling | Sourmash | 4.7.0 | GTDB R214 and Genbank 2022-04 (for Fig. S1 only) | | gather -p k=31,scaled=1000,abund |
|  | GTDB-Tk | 2.2.5 | GTDB R214 | | classify_wf |
| Phylogeny | PhyloPhlAn | 3.0.3 | SGB.Oct22 | | --diversity high --accurate |
| Dark matter exploration | BLAST+ | 2.12.0 | 2022_03_23 nt | | -evalue 0.01 -qcov_hsp_perc 75 -word_size 20 -max_target_seqs 5 |
|  | seqtk | 1.3 |  | |  |
| R – Statistical analyses | vegan | 2.6-4 |  | |  |
|  | R | 4.2.1 |  | |  |
|  | DESeq2 | 1.42 |  | |  |
|  | phyloseq | 1.46.0 |  | |  |
|  | DHARMa | 0.4.6 |  | |  |
|  | lme4 | 1.1-35.1 |  | |  |
|  | ANCOMBC | 2.4.0 |  | |  |
|  | ComplexHeathmap | 2.18.0 |  | |  |
| R – Figures | ggplot2 | 3.4.4 |  | |  |
|  | ggtree | 3.10.0 |  |  | |
|  | ggtreeExtra | 1.12.0 |  |  | |
|  | ggeffects | 1.4.0 |  |  | |
|  | treeio | 1.26.0 |  | |  |
|  | patchwork | 1.2.0 |  | |  |
|  | cowplot | 1.1.3 |  | |  |
|  | ape | 5.7-1 |  | |  |
|  | metBrewer | 0.2.0 |  | |  |
|  | ggnewscale | 0.4.10 |  | |  |
|  | scales | 1.3.0 |  | |  |
| R – Analysis and statistics | tidyverse | 2.0.0 |  | |  |
|  | bestNormalize | 1.9.1 |  | |  |
|  | betareg | 3.1-4 |  | |  |
|  | emmeans | 1.10.1 |  | |  |
|  | rstatix | 0.7.2 |  | |  |
|  | taxize | 0.9.100 |  | |  |
|  | permute | 0.9-7 |  | |  |
| R – Parallel processing | doParallel | 1.0.17 |  | |  |
|  | foreach | 1.5.2 |  | |  |

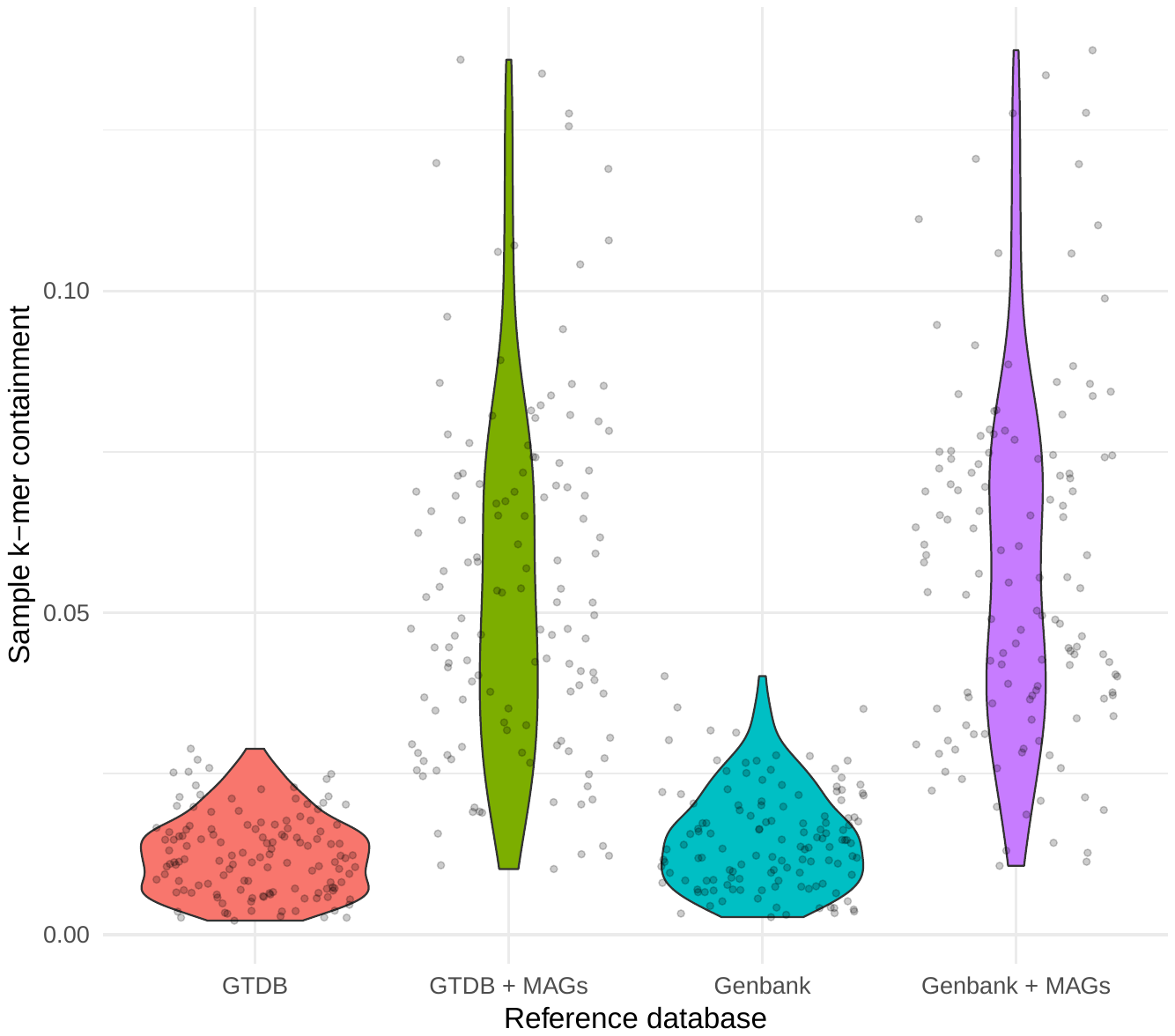

Supplementary Figure S1. Comparison of sample k-mer containment using the GTDB R214 and GenBank 2022-03 reference databases, with and without the inclusion of metagenome-assembled genomes identified as novel (nMAGs). The violin plots illustrate the estimated increase in sample coverage when nMAGs are incorporated into each reference database. Default GTDB and GenBank databases serve as baselines, with enhancements observed upon the addition of nMAGs.

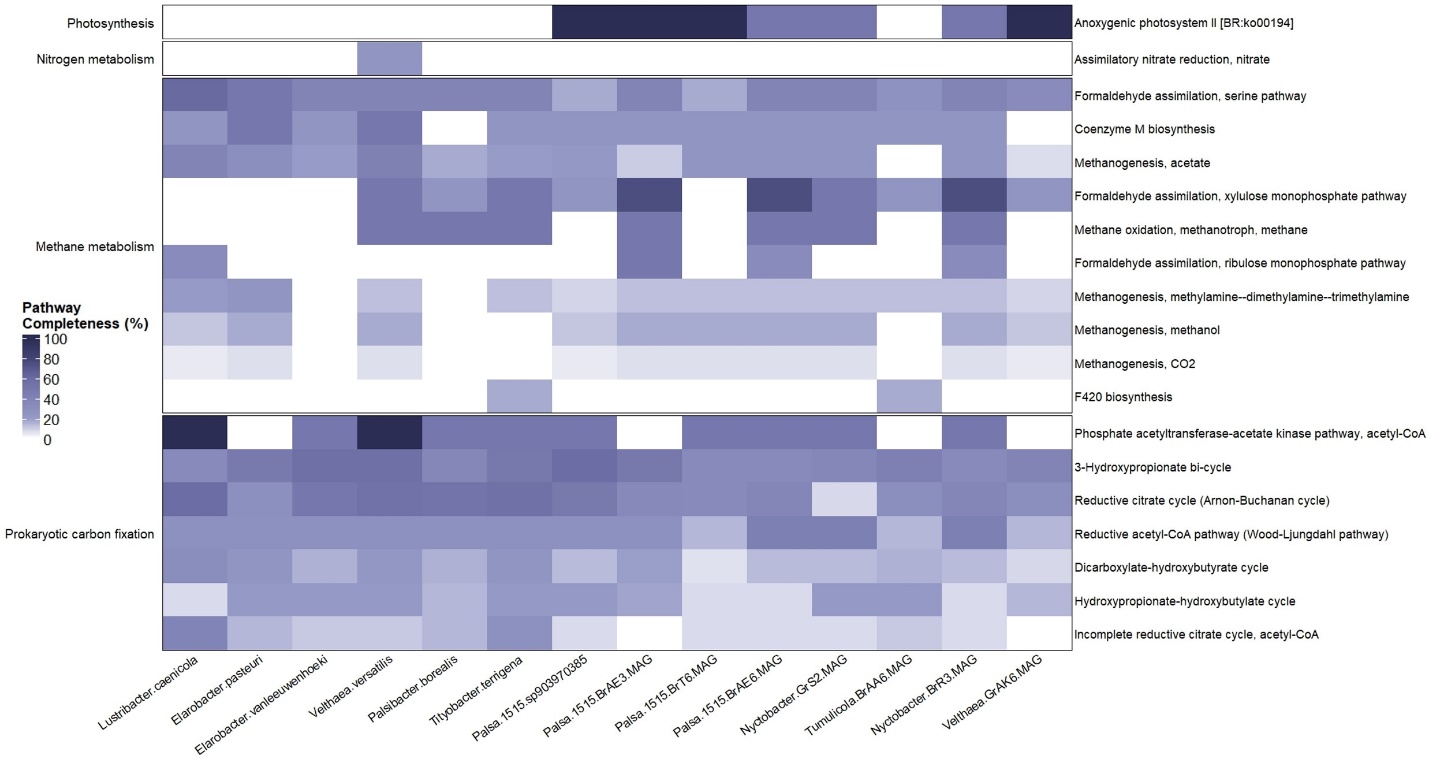

Supplementary Figure S2. Potential metabolic capacity of every Eremiobacterota detected across samples. The pathway completeness (%) for four pathway groups (photosynthesis, nitrogen and methane metabolism, and prokaryotic carbon fixation) is indicated for each genome (blue scale).

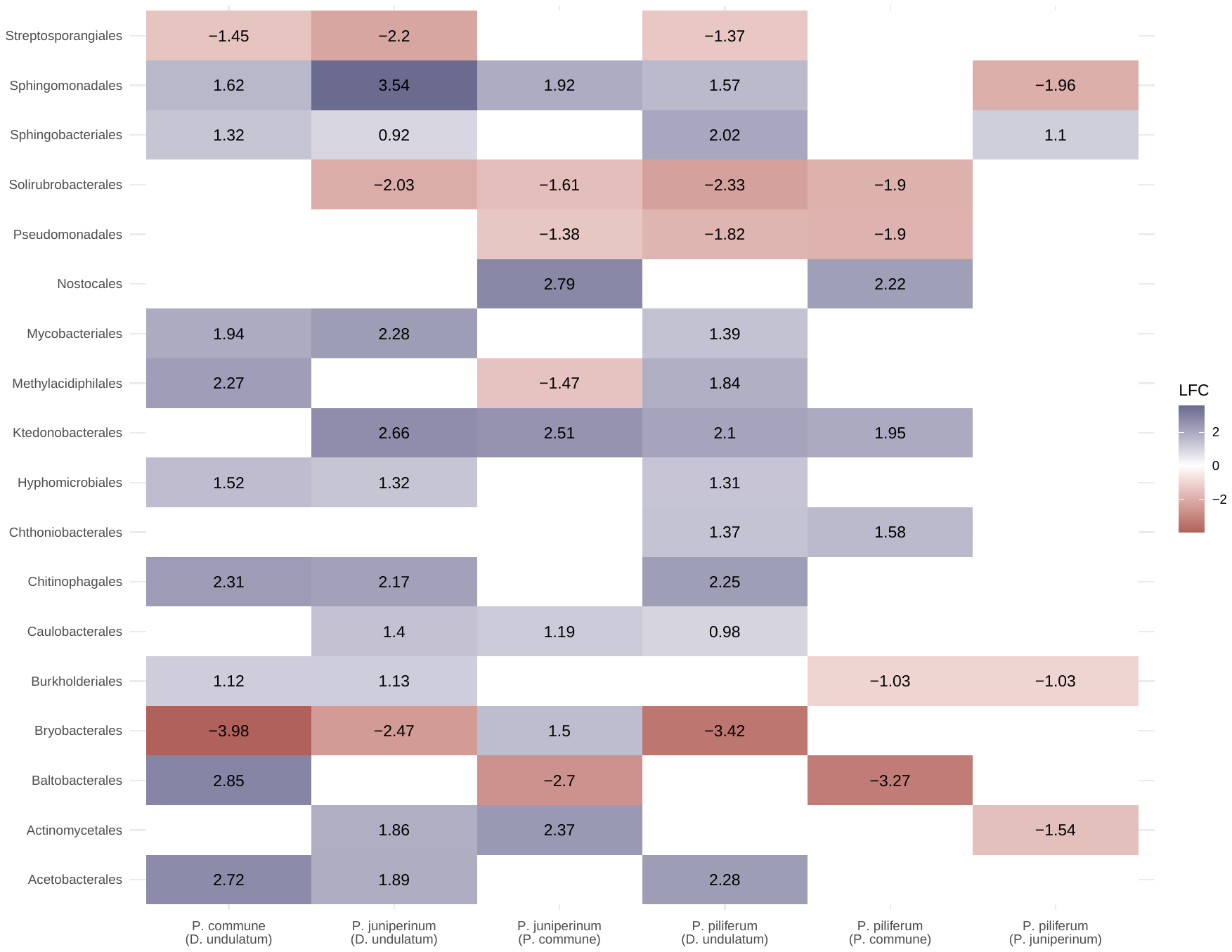

Supplementary Figure S3. Differentially abundant orders in moss species. LFC: log2-fold change in absolute abundances. All values shown are significantly differentially abundant (p <0.01). Positive LFC (blue) indicates increase in moss species not in parentheses relative to moss species in parentheses, whereas negative LFC (red) indicates increase in the opposite. Blanks indicate lack of significant differential abundance (LFC not reported).

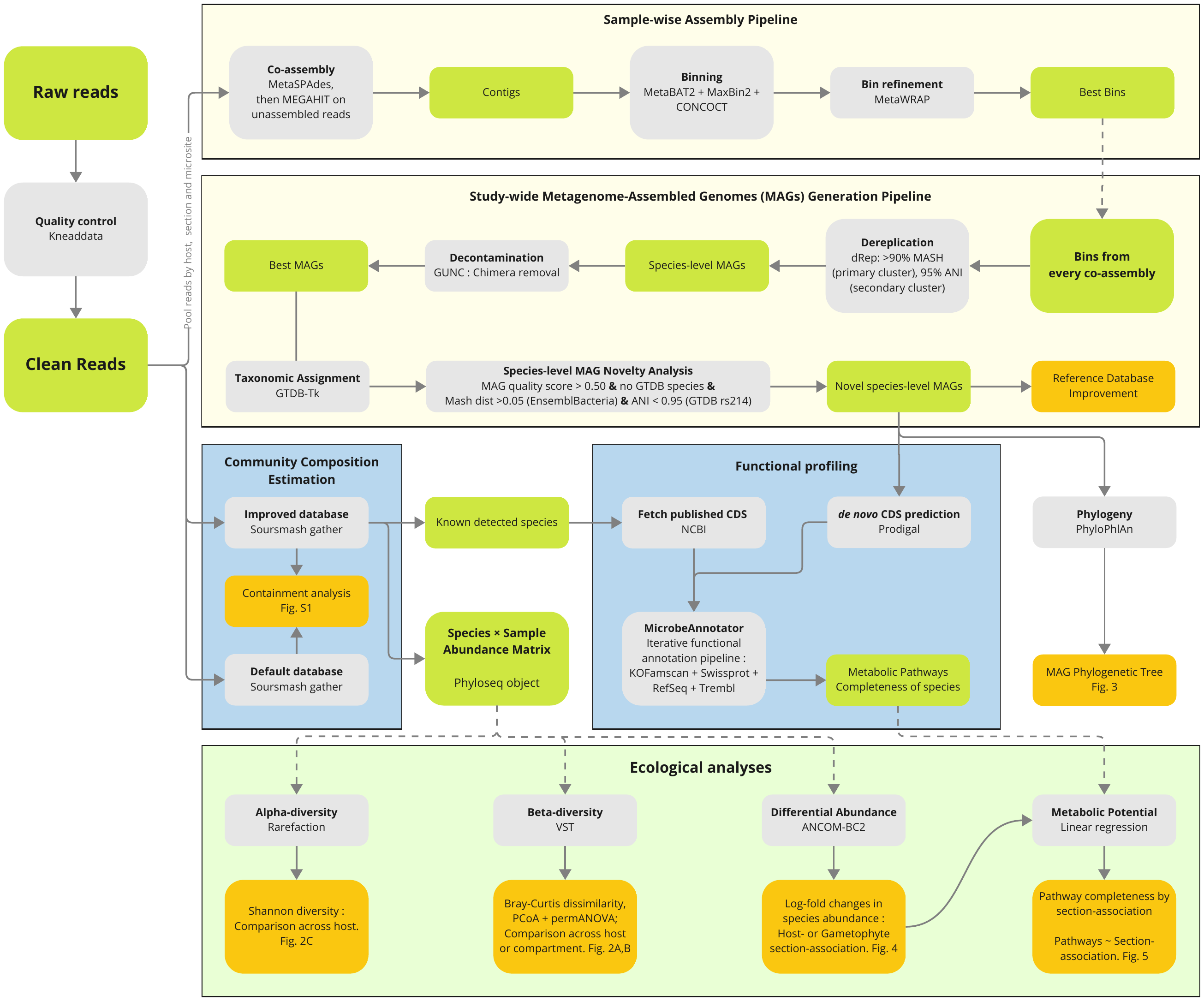

Supplementary Figure S4. Bioinformatic metagenomics pipeline. Boxes colored in grey represent processes (or programs); in green, input/output data; in orange, analyses and results.

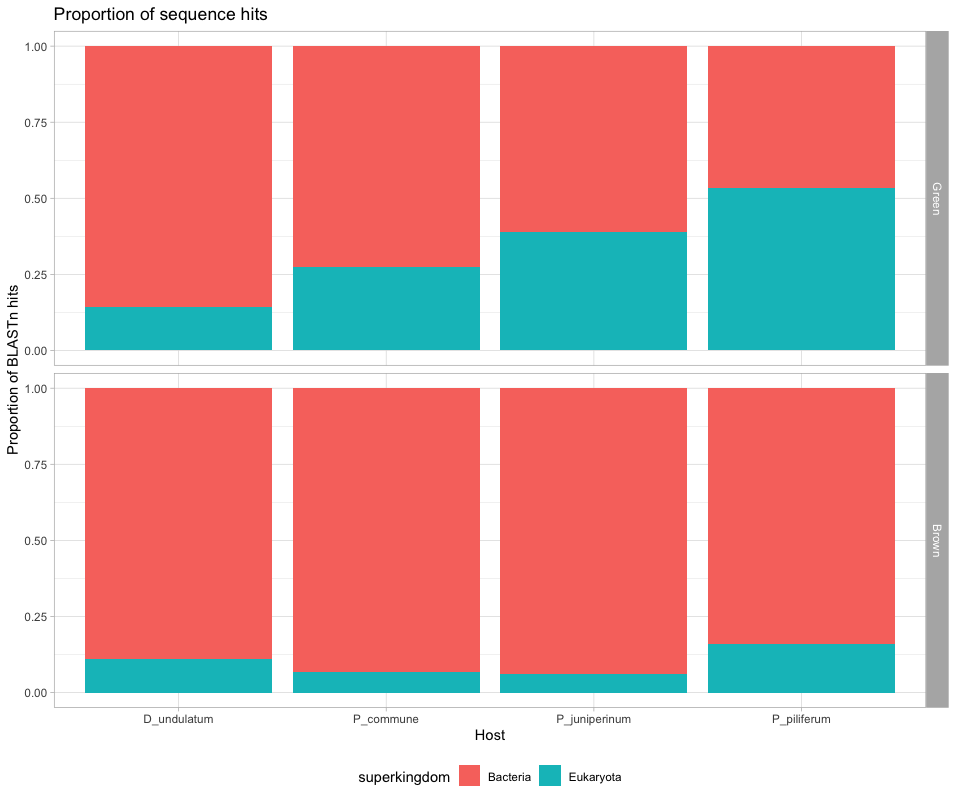

Supplementary Figure S5. Hits of sequences at the kingdom level across moss species and gametophyte sections using BLASTn. Proportions of 100,000 randomly selected reads per sample, categorized into Bacteria (red) and non-microbial Eukaryota (cyan) superkingdoms. The data indicates a predominant bacterial presence in the 5 top BLASTn sequence hits, alongside a notable proportion of eukaryotic non-microbial reads.

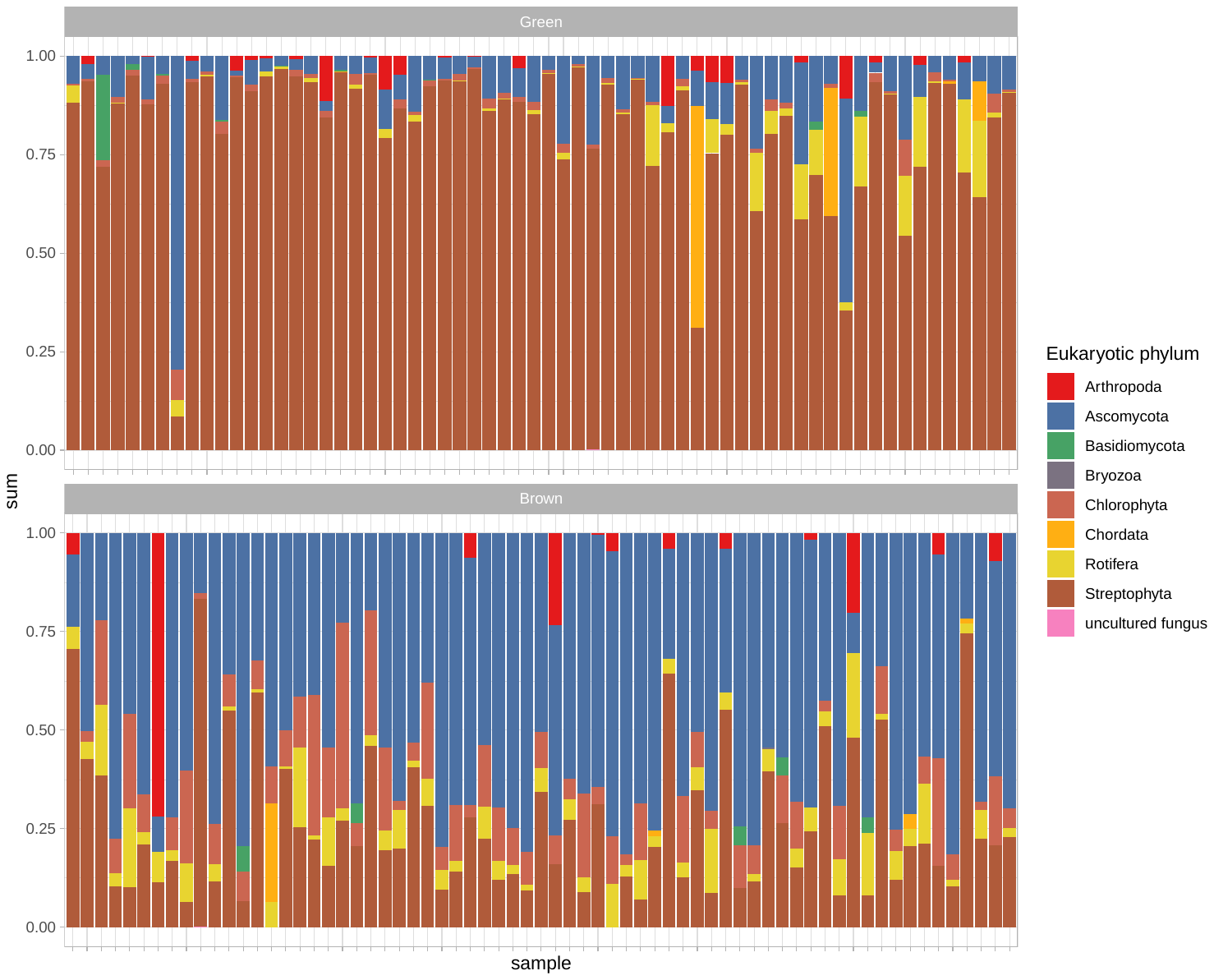

Supplementary Figure S6. Proportion of eukaryotic phyla sequence hits using BLASTn. 100,000 randomly selected reads per sample were used, results are shown for the green (top panel) and brown (bottom panel) gametophyte sections.
